# The quorum-sensing lexicon of *Salmonella* ameliorates acid stress in the host by a non-canonical mechanism

**DOI:** 10.1101/2025.03.26.645541

**Authors:** Anmol Singh, Abhilash Vijay Nair, Shashanka Aroli, Suman Das, Subhrajit Karmakar, R. S. Rajmani, Santanu Mukherjee, Umesh Varshney, Dipshikha Chakravortty

**Affiliations:** Department of Microbiology and Cell Biology, Division of Biological Sciences, Indian Institute of Science, Bengaluru, India; Department of Organic Chemistry, Indian Institute of Science, Bengaluru, India; Molecular Biophysics Unit, Indian Institute of Science, Bengaluru, India; Adjunct Faculty, School of Biology, Indian Institute of Science Education and Research, Thiruvananthapuram, India

**Keywords:** Autoinducer-2, acidic pH, macrophage, PhoP/PhoQ, *Salmonella* pathogenicity island-2

## Abstract

The intestinal milieu is largely characterized by the complex array of chemical compounds produced through the metabolic activity of resident microbiota. Enteric pathogens like *Salmonella*, which have evolved refined mechanisms to persist within this environment, utilize these microbial metabolites and self-produce quorum molecules as molecular cues to identify ecological niches and modulate their survival and virulence strategies. *Salmonella* quorum sensing involves producing and detecting Autoinducer-2 (AI-2) signaling molecules. Our research reveals that *Salmonella* Typhimurium enhances AI-2 biosynthesis and transport under acidic conditions, aiding environmental adaptation and facilitating pathogenesis in macrophages. AI-2 signaling regulates the pH-sensing two-component system genes, *phoP/phoQ*, ensuring cytosolic pH homeostasis, survival, and acid tolerance. It also involves regulating the lysine/cadaverine-mediated acid tolerance response and maintaining bacterial cytosolic pH. Furthermore, we show that the repressor LsrR protein, apart from the *lsr* promoter, binds to only one strand of the *phoP* promoter via its Y25 and R43 amino acids to negatively modulate *phoP* expression. Additionally this signaling ameliorates the intracellular survival by modulating *Salmonella* Pathogenicity Island-2 (SPI-2) regulators (*ssrA/ssrB*) and SPI-2 effector expression via PhoP. Mouse models demonstrate that AI-2 signaling is essential for colonizing the primary and secondary infection sites, showcasing its critical role in pathogenesis and low pH survival mechanisms in *Salmonella*.

## Introduction

*Salmonella* is considered a major foodborne enteric pathogen worldwide (1), and it gains access to the host through contaminated food and water. The bacteria stealthily survive the acidic pH of the stomach and the hostile environment of the host macrophages. Within the host cells, *Salmonella* resides in the *Salmonella-*containing vacuole (SCV), a membrane-bound compartment that provides with an acidic niche facilitating utilization of the SPI-2 encoded genes for the intracellular proliferation of the bacterium (2-4). *Salmonella enterica* serovars infection in humans can elicit a wide range of clinical outcomes, spanning asymptomatic carriage, acute self-limiting gastroenteritis, invasive systemic disease with associated bacteremia, and enteric (typhoid) fever(5).

The community behaviour of bacteria, known as quorum sensing, protects them from various stress factors such as exposure to ultraviolet light, acids, detergents, or antimicrobial agents (6, 7). Also, AI-2 signaling is linked to bacterial survival and pathogenesis, for example, in *Lactobacillus sp.,* AI-2 helps in *s*urvival under acidic conditions (8, 9). *Salmonella* recognizes a chemically distinct quorum sensing signal called autoinducer 2 (AI-2), synthesized by the enzyme LuxS (10), which is maximally produced during its exponential growth phase (11). As bacterial quorum increases, extracellular AI-2 accumulates and, upon reaching a threshold, it is sensed by LsrB protein in *Salmonella*. AI-2 is then transported into the cytosol by an ABC transporter (LsrA, LsrB, LsrC, and LsrD) encoded by the *lsr* operon and phosphorylated by the LsrK kinase. The phosphorylated AI-2 binds LsrR repressor protein, and relieves the LsrR mediated repression of the *lsr* operon to enable further import of AI-2 (12, 13).

The LuxS/AI-2 signaling regulates the expression of oxidative stress response related genes such as *sodA*, *sodCI*, and *sodCII*, which play an essential role in survival of *Salmonella* within macrophages (14). Further, the release of LsrR from DNA by AI-2 is essential for SPI-1 transcription and flagella expression in *Salmonella* (15, 16). However, the molecular basis of this remains unclear, not only because of the diverse and distinct survival strategies of *Salmonella* but also due to the limited understanding of the AI-2 regulatory mechanism and its role in enhancing pathogenesis beyond the *lsr* operon.

As an enteric pathogen, *Salmonella* has to thrive in diverse and competitive environments in the gastrointestinal tract. It needs to sense and adapt to a variety of environmental barriers such as host physiology, nutrient availability, osmotic stress, and competition with other gut bacteria(17). Counteracting and conquering the acidic pH of the stomach and acidic intravacuolar environment is an indispensable survival strategy for *Salmonella.* We contemplated that the interbacterial communication and signaling systems might control the pH sensing and tolerance responses. Here, we show that LuxS/AI-2 signaling regulates a two-component system *phoP*/*phoQ* to activate the acid tolerance system in *Salmonella* and tune the SPI-2 encoded genes, enhancing its survival in *in vitro* and *in vivo* models. We found that LsrR, the regulator of LuxS/AI-2 signaling, binds to the *phoP/phoQ* promoter via its Y25 and R43 amino acid residues, apart from its known regulation of the *lsr* operon. This way, LuxS/AI-2 coordination facilitates dynamic changes of the *Salmonella* transcriptome according to the surrounding environment and promotes rapid adaptation to different environmental niches to increase population fitness.

## Materials and Methods

### Bacterial strains and Growth conditions-

The wild-type *Salmonella enterica* serovar Typhimurium strain 14028S (STM WT) used in all experiments in this study was a kind gift from Professor Michael Hensel of the Max Von Pettenkofer-Institute for Hygiene und Medizinische Mikrobiologie in Germany. The bacterial strains were revived on LB agar with or without antibiotics. The LB broth culture of wild-type, knockout, and complemented strains was cultured at 37°C (170 rpm) in an orbital shaker incubator. Antibiotics Kanamycin (50μg/ml), Chloramphenicol (25μg/ml), and Ampicillin (50μg/ml) were used wherever required. *Vibrio campbellii* ATTC BAA-1117 strain was used for autoinducer (AI-2) bioassay. This strain was grown in ATCC Medium: 2746 Autoinducer Bioassay (AB) Medium.

A growth kinetics study was conducted in LB and M9 minimal media (47) (48) (**Table S2**). Briefly, STM WT and mutants were inoculated in fresh LB broth (1:100 ratio from overnight primary culture). The cultures were incubated at 37°C (170 rpm) in an orbital shaker incubator. The OD at 600nm was measured at regular intervals until 24 hours post-inoculation.

### Bacterial gene knockout and strain generation

*Salmonella enterica* serovar Typhimurium strain 14028S and isogenic mutants were used for all assays. *luxS*, *lsrB*, *lsrK,* and *lsrR* gene knockout strains were made by *λ*-Red recombination system as described previously in one gene step chromosomal gene inactivation method demonstrated by Datsenko and Wanner (49). Briefly, STM WT strains were transformed with pKD46 plasmid expressing λ-red recombinase system under Arabinose inducible plasmid. The pKD3 and pKD4 plasmid were used as the template to amplify the chloramphenicol and kanamycin resistance cassette with knockout primers **(Table S3).** The amplified reaction products were purified using chloroform isopropanol precipitation. The purified PCR product was transformed into pKD46-containing STM WT expressing λ-Red recombinase system. The transformed cells were plated on LB agar with chloramphenicol and kanamycin accordingly. The knockout strains were confirmed by PCR using primers **(Table S3)** corresponding to the regions ∼100bp upstream and downstream of the relevant genes.

For complement strain, colony PCR amplified the *luxS* gene with gene-specific primers (containing BamH1 and HindIII restriction site sequence) (Table S3). The amplified PCR product was purified by chloroform isopropanol precipitation and cloned into the empty pQE60 vector using BamH1 and HindIII sites. The double-digested insert and vector were subjected to ligation by T4DNA ligase in ligation buffer (NEB) overnight at 16°C. The respective ligated vector was transformed into the STM Δ*luxS* strains to generate the complemented strain. Complement strain was initially confirmed by colony PCR with cloning primers and internal primers of the genes, and finally by restriction digestion of recombinant plasmid isolated from complement strains to study insert release.

### Cell culture and maintenance

Murine macrophage cell line RAW 264.7 was maintained in Dulbecco’s Modified Eagle Medium (DMEM, Lonza), supplemented with 10% FBS (Gibco) and 1% penicillin-streptomycin (Sigma-Aldrich) in a humidified incubator with 5% CO2 at 37 °C. The cells were seeded into the respective cell culture plate for each experiment to conduct intracellular survival assays and intracellular gene expression experiments.

### Peritoneal macrophage isolation

The primary macrophage isolation was performed as previously described (50). Briefly, C57BL/6J mice (5-6 weeks old) were injected with Brewer’s thio-glycolate medium (8% w/v, HIMEDIA). At four days of treatment, 5 ml of cold PBS was injected into the peritoneal cavity, followed by aspiration of the fluid from the cavity and dispensed into a 15 ml centrifuge tube. The centrifugation was done at 300g for 10 min, and the resuspended cell pellet [in Roswell Park Memorial Institute 1640 (RPMI 1640)] was supplemented with 10% FBS and an antibiotic cocktail (penicillin-streptomycin). Cells were counted by hemocytometer and seeded onto 24 well cell culture plates.

### Chemical synthesis of acetonide-protected (*S*)-4,5-dihydroxy-2,3-petanedione (DPD)

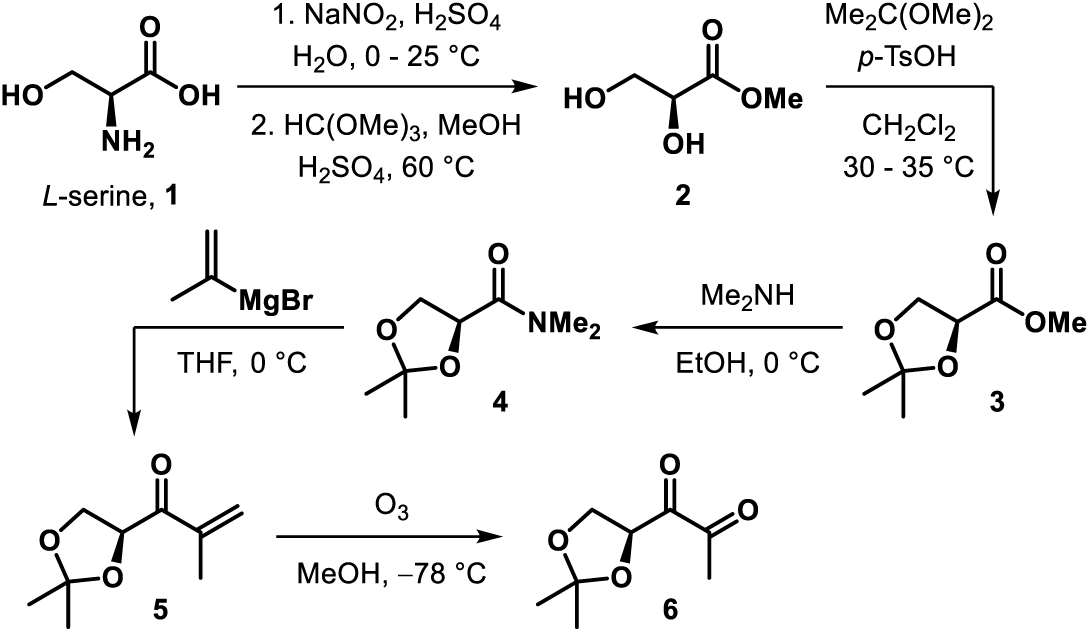

Acetonide-protected DPD **6** was synthesized from commercially available *L*-serine **1** in five steps (for details, see the supplemetary Information). These steps include stereoretentive displacement of the amino group of *L*-serine through diazotisation and esterification to form methyl (*S*)-2,3-dihydroxypropanoate **2** (51) followed by protection of 1,2-diol as acetonide (**3**(52). Transformation of methyl ester of **3** to *N*,*N*-dimethyl amide **4** (53) paved the way for its conversion to isopropenyl ketone **5** by reacting with isopropenyl Grignard reagent. Ozonolytic cleavage of the alkene in **5** resulted in acetonide-protected (*S*)-DPD **6**(54).

### AI-2 bioassay

The culture of *Salmonella* Typhimurium WT and mutants was obtained at 0h, 1h, 3h, 6h, 9h, and 12h, and centrifuged at 6000rpm for 10 min. After filtration using a 0.22 µm filter, the supernatant was stored at -20 °C until required. The AI-2 was quantified using the sensor strain *Vibrio campbellii* ATTC BAA-1117 (strain designation *V. harveyi* BB-170, which only responds to AI-2). The AI-2 bioassay was performed as previously described (55) (6) with minor modifications. A single colony of *V. harveyi* BB-170 was incubated in 5 ml of Autoinducer Bioassay medium (ATCC Medium: 2746 AB Medium) at 30°C and 200 rpm. A 1:1000 dilution was obtained by transferring a 10 µl sensor strain culture to 10 mL of fresh AB media. *Salmonella* cell-free culture supernatant (20 μl) was incubated with a 180 ml diluted sensor strain. The mixture was added to a white, flat-bottomed 96-well microtiter plate (Corning, USA), followed by shaking in a rotary shaker at 200 rpm, at 30°C. Luminescence was measured at 3 h in a fluorescent microtiter plate reader. Autoinducer signal calculated as – (luminescence produced by *Vibrio* in the presence of *Salmonella* Typhimurium free supernatant - luminescence in media control)/ luminescence produced by *Vibrio* with sterile media. To validate the activity of exogenous DPD molecule and BF-8 inhibitor, an autoinducer bioassay was performed in the presence of DPD and BF-8 inhibitors (Cat# 53796, purchased from Sigma).

### Gentamicin protection assay

RAW 264.7 macrophages and peritoneal macrophages isolated from C57BL/6J mice were seeded in tissue culture plates for infection. Cells were infected with STM WT, STM Δ*luxS,* STM Δ*lsrB,* STM Δ*lsrK,* STM Δ*lsrR,* and *STM*Δ*luxS:luxS* (stationary phase culture growing overnight in LB broth). The Multiplicity of Infection (MOI) of 10 was used for the intracellular survival assay. MOI of 20 was used for the *Salmonella* Typhimurium gene expression study (qRT-PCR). Following the infection of STM strains into the cell line, the plate was spun for 5 min at 600 rpm using a Rota-Superspin R–NV swing bucket centrifuge. The plate was then incubated for 25 min at 37 °C with 5% CO2 in a humidified incubator. The infected cells were washed twice with 1X PBS and treated with a 100 μg/ mL concentration of gentamycin. After 1 h, the media were changed with a reduced concentration of gentamycin (25 μg/ mL) and incubated until the desired time point. The time points of 2h, 6h, and 16h were taken for the qRT-PCR samples and 2h and 16h for the intracellular survival assay.

Also, for studying the effect of exogenous DPD in the recovery of the phenotype of mutant *luxS,* were grown *in vitro* with supplementation of 25 μM, 50 μM,100 μM, and 200 μM DPD molecule and 10%, 20%, 30%, and 50% spent media of STM WT collected at 3h in LB growing. For inhibition, the inhibitor (Z)-4-bromo-5-( bromomethyllene)-3-methylfran-2(5H) known as BF-8 treatment, was added to STM WT in LB media. DPD or spent media supplemented and BF-8 treated culture was used for gentamycin protection assay.

### Intracellular survival assay and phagocytosis assay

The cells were lysed with 0.1% triton-X 100 in PBS at specific time points post-infection. For the intracellular survival assay, 2h and 16h post-infection samples were collected and plated, and the corresponding CFU at 2h and 16h was determined. Fold proliferation was determined using the formula [ CFU at 16 h]/ [ CFU at 2 h]. For phagocytosis, the bacterial number was determined in the inoculum and 2h time point post-infection. Percent phagocytosis was by using the formula- (CFU at 2 h)/ (CFU of pre-inoculum)]×100.

### Confocal microscopy

To study the *Salmonella* proliferation and their intracellular vacuolar status, all STM strains (STM WT, STM Δ*luxS,* STM Δ*lsrB,* STM Δ*lsrK,* and STM Δ*lsrR*) were transformed with pFPV 25.1-mCherry-Amp^R^ plasmid, and used to infect RAW264.7 cells. Briefly, 1×10^5^ RAW264.7 cells were seeded on the coverslip and infected with different strains of bacteria at MOI 25. After appropriate post-infection durations, cells were washed twice with PBS and fixed with 3.5% paraformaldehyde (PFA), and incubated with a specific antibody (anti-mouse LAMP-1) in a blocking buffer containing 2% BSA and 0.01% saponin for 2 hours at RT or overnight at 4℃. The cells were washed twice with PBS and incubated with appropriate secondary antibody tagged with fluorochrome for 1 hour at RT. The coverslips were then mounted onto a clean glass slide and imaged under a confocal microscope (Zeiss LSM-710) using a 63X objective.

### *In vitro* Cell culture competition Assay

RAW 264.7 macrophages were infected at a 1:1 ratio with STM WT and STM Δ*luxS.* At 16 h, post-infection cells were lysed, and the sample was collected as described above. The competition was also done with the coculture of STM WT and STM Δ*luxS*. This involved the co-cultivation of STM WT and STM Δ*luxS* together, followed by infection in RAW264.7 macrophages. The competitive index was calculated for a WT strain and Δ*luxS* by dividing the ratio between CFU (STM Δ*luxS*) and CFU (STM WT) by the ratio of both strains in the inoculum.

### RNA isolation and RT-qPCR

For gene expression study in Luria Bertani (LB) media and F-media (acidic media that mimics the cell vacuole environment) (56) (48), overnight-grown culture in LB media was subjected to subculture (at 1:100) in LB or F-media (**Table S1**). At 3h, 6h,9h, and 12h, bacterial cell pellets were resuspended in TRIzol (from TaKaRa, RNA isoPlus-9109) and kept at -80°C. Total RNA was isolated by chloroform extraction followed by isopropanol precipitation. To evaluate the quality, the amount of RNA was quantified in nanodrop (ThermoFischer) and examined on a 2% agarose gel.

To produce cDNA, 3 μg of RNA sample was treated with DNase I (TaKaRa) at 37°C for 1 h followed by heating for 10 min at 65°C. DNA-free RNA samples were used to make cDNA using the PrimeScript RT reagent Kit provided by TaKaRa (Cat# RR037A).

For gene expression in *Salmonella* Typhimurium upon infection in RAW 264.7 macrophages, infections were performed at MOI of 20. At 2h, 6h, and 16h post-infection, the cells were lysed using TRIzol and kept at -80°C. Total RNA isolation and cDNA synthesis were carried out using the manufacturer’s protocol. The list of expression primers is provided in **Table S3**.

### Acid survival assay

Overnight cultures of STM WT, STM Δ*luxS,* STM Δ*lsrB,* STM Δ*lsrK,* and STM Δ*lsrR* were adjusted to OD600 - 0.3. For survival in 1XPBS, pH was adjusted to 3,4,5,6,7, or 8 using concentrated HCl. Strains of *Salmonella* Typhimurium (10 ^7^ bacteria/ml) were exposed to different pH as mentioned in 1X PBS and incubated for 2h. At 2h post-treatment, CFU were enumerated by plating on LB agar media. The same protocol was followed for survival in LB media with similar pH ranges. For the strain of STM harboring plasmids pQE60 complemented with *phoP*/*phoQ*, a survival assay was done in pH of 3, 4, and 5 in 1X PBS.

### Acid tolerance assay

A standard acid tolerance assay was performed as previously described with slight modification (22). Briefly, acid tolerance assay (ATR) was conducted with strains grown overnight at 37°C in LB broth containing the appropriate antibiotic. The culture corresponding to 0.3 OD600 of the overnight culture (unadapted) was centrifuged and the pellet was resuspended into 2 ml of LB broth (pH 5) and incubated at 37°C with shaking for 2 h (known as adapted culture). The acid challenge of unadapted and adapted cultures involved readjusting the pH to 3. CFU was determined for adapted and unadapted cultures after exposure to pH 3 (2 h of pH 3 treatment). ATR assay was also done with the strain of *Salmonella* containing plasmid pQE60 complemented with *phoP/phoQ* gene.

### Intracellular pH measurement by pHuji plasmid

The plasmid pBAD:pHuji (Plasmid #61555) was used to transform STM WT, STM Δ*luxS*, STM Δ*lsrB*, STM Δ*lsrK*, and STM Δ*lsrR*. pHuji is a pH-sensitive red fluorescent protein demonstrating a more than 20-fold fluorescent intensity change from pH 5.5 to 7.5 (57). Overnight-grown bacterial strains were subjected to a pH range of 3.0 to 8.0 in phosphate buffer (PBS) in 40µM of sodium benzoate to ensure equilibration to the desired pH. The resulting fluorescence intensity ratio, obtained from flow cytometric analysis, was plotted as a function of pH and fitted to get a standard curve. The standard curve was used to interpolate ratios measured within the intracellular pH of bacteria in LB media, F-media, and RAW 264.7 macrophages.

### Mass-spectrometry for the determination of lysine and cadaverine

Overnight grown STM WT, STM Δ*luxS*, STM Δ*lsrK,* STM Δ*lsrB,* and STM Δ*lsrR* were sub-cultured in acidic pH 5 LB media and incubated at 37℃ until mid-log phase. Thereafter, bacteria were spun down and the spent media were collected. Spent media were kept with an equal volume of acetone at -20℃ for overnight. Next, samples were collected after centrifugation at 13k rpm for 10 min. Samples were analysed in Orbitrap fusion connected to UHPLC Vanquish (Thermo Scientific). Water and acetonitrile with 0.1% formic acid were used as a mobile phase. The flow rate was kept at 0.3ml/min linear gradient was started with 5% to 95% using Hypersil Gold C18 column (2.1X100mm & particle size 1.9µ).

### Plasmid construction, and site-directed mutagenesis

The vector pET28a(+) was used to construct plasmids p*lsrR* for the overexpression of LsrR as a C-terminal 6xHis-tagged fusion protein. The DNA fragment containing *the lsrR* coding region was amplified by polymerase chain reaction (PCR) from the chromosome of *Salmonella* Typhimurium 14028s using primers (F-lsrR and R-lsrR containing restriction site NcoI and XhoI, respectively). The PCR products were digested with NcoI and XhoI, and ligated with NcoI and XhoI digested pET28a(+). The inserts were checked by DNA sequencing.

For Y25A and R43A site-directed mutagenesis in LsrR protein, PCR for SDM was set up with template DNA from pETa(+)- *lsrR* and amplified with Q5 polymerase (NEB). The PCR products were digested with DpnI and transformed into *E. coli* DH5-alpha. Y25A and R43A double mutants were generated using pET28a(+) *lsrR* R43A plasmid as a template for SDM using primers specific to the second set of mutations. All plasmids were confirmed by DNA sequencing.

### Overexpression, and purification of LsrR

pET28a(+)-*lsrR* and its mutants containing C-terminal 6xHis tag were transformed into *E. coli* Rosetta (DE3). The transformed colonies were inoculated into 5 ml LB containing kanamycin (Kan) and chloramphenicol (Cm) and grown overnight. Inoculum (1%) was added to 1.2 L LB containing Kan and Cm, grown at 37°C to an OD600 of 0.4 under shaking, and thereafter supplemented with 0.1 mM IPTG, and grown at 16℃ overnight. Cells were harvested by centrifugation at 4℃, resuspended in 10 ml buffer A [20 mM Tris–HCl pH 8, M NaCl, 10% glycerol (v/v), 1.5 mM β-mercaptoethanol, 1 mM PMSF] lysed by sonication, and centrifuged at 4℃ in a pre-cooled centrifuge at 10000 rpm for 10 min. The supernatant was loaded onto a 1 ml Ni-NTA column equilibrated with buffer A, washed with 20 ml buffer A, and eluted with a gradient of imidazole (20–500 mM) in the same buffer. The fractions were analysed on 12% SDS-PAGE. Fractions enriched for LsrR were pooled, loaded onto a Superdex75 gel filtration column, and eluted in buffer B [20 mM Tris–HCl pH 8, 1M NaCl, 10% glycerol (v/v) and 1.5 mM β-mercaptoethanol]. The purity of LsrR was checked on 12% SDS-PAGE **(Fig. S7H)**. Fractions with apparent homogeneity were pooled, concentrated using a 10 kDa cut-off Centricon (Millipore), and estimated by Bradford’s method using bovine serum albumin (BSA) as standard. The proteins were dialyzed against buffer A containing 50% glycerol (v/v) and stored at –20°C.

### Gel shift assay

HPLC-purified 90bp single-strand DNA oligos of the *lsrR* and *phoP* operon promoters (Sigma) were used for a gel shift assay. Binding reactions involved incubating promoter DNA with varying concentrations of LsrR or its mutated form. The incubation buffer contained 50 mM Tris-Cl (pH 7.5), 150 mM NaCl, 3 mM magnesium acetate, 0.1 mM EDTA, and 0.1 mM DTT. After 15 min at room temperature, the mixture was combined with gel loading buffer (60% 0.25× TBE, 40% glycerol, 0.2% bromophenol blue) and run on a 6% native polyacrylamide gel. The staining was done for DNA with SYBR™ Gold Nucleic Acid Gel Stain (Invitrogen) and analyzed as per the manufacturer’s instructions.

### *In vivo* animal experiment

Oral gavaging of 10^7^ CFU of STM WT, STM*ΔluxS*, STM*ΔlsrB*, STM*ΔlsrK*, STM*ΔlsrR*, and STM*ΔluxS:luxS* was used to infect 5- to 6-week-old C57BL/6J mice. Post five days of infection, the intestine (Peyer’s patches), MLN (mesenteric lymph node), spleen, liver, and blood were aseptically extracted (in a Biosafety level 2 cabinet) to examine the colonization in the organs. The CFU were enumerated on differential and selective *Salmonella*-*Shigella* (SS) agar.

### Mice survival assay

Male C57BL/6, aged 5 to 6 weeks, were obtained from the Central Animal Facility, IISc. The mice were given 10^8^ CFU of each strain (overnight grown cultures) orally to compare the survival and weight alterations of mice post-infection. Following infection, the mice were observed every day to determine their survival and weight, and the results were expressed as a percentage of survival (Kaplan-meier curve) and weight reduction.

All experiments comply with the rules set forth by the Indian Institute of Science, Bangalore’s IEAC. The approved protocol number is CAF/ Ethics/852/2021. The Institutional Animal Ethics Committee approved every animal experiment, and the National Animal Care Guidelines were scrupulously followed.

### Statistical analysis

As mentioned in figure legends, each experiment has been independently repeated two to five times. GraphPad Prism 8.4.3 was utilized for all statistical analyses. As the figure legends state, the statistical analyses included an unpaired, two-tailed Student’s t-test, One-way ANOVA with Dunnet’s post hoc test, and Two-way ANOVA with Tukey’s post hoc test. A non-parametric one-way ANOVA (Kruskal Wallis) test with Dunn’s post-hoc test was performed for the animal experiment. P-values less than 0.05 were regarded as significant. The analysis is presented as mean ±SD or mean ±SEM with information on group sizes and p values mentioned in the respective figure legends.

## Results

### LuxS/AI-2 quorum sensing is required by *Salmonella* Typhimurium to survive in the macrophages

We first investigated the levels of AI-2 produced at different growth phases from *Salmonella* Typhimurium (STM) WT grown at 37℃ in LB media by using the *Vibrio harveyi* BB-170 reporter strain (which produces luminescence responses to AI-2). Consistent with previous studies (16), AI-2 dependent light production by *V. harveyi* BB-170 is maximum in the spent medium of the mid-log phase culture of STM, and it gradually decreases towards the late stationary phase **(Fig. 1A; Fig. S1A)**. Also, while *Salmonella* upregulated the expression of the *lsr* operon genes (*lsrB*, *lsrK*, *lsrR*) during its mid-log phase to early stationary phase of growth **(Fig. 1B)**, it did not alter the expression of *luxS* during this period **(Fig. 1C)**. And the deletion of *luxS* and *lsr* operon genes (*lsrK*, *lsrB*, *lsrR*) in STM did not affect the *in vitro* growth in LB and minimal media **(Fig. S1B, C)**. Next, we observed that in comparison to the STM WT, the quorum-sensing mutants STM*ΔluxS*, STM*ΔlsrB*, and STM*ΔlsrK* displayed increased phagocytosis in RAW 264.7 macrophages **(Fig.1D)**. However, both STM*ΔlsrR*, and *luxS* complemented STM*ΔluxS* behaved same as STM WT **(Fig.1D)**. Importantly, the STM*ΔluxS*, STM*ΔlsrB*, and STM*ΔlsrK* but not the STM*ΔlsrR*, and *luxS* complemented STM*ΔluxS* showed a decreased ability to proliferate in RAW 264.7 cells **(Fig. 1E; Fig. S1E, F)**.

**Fig. 1:**
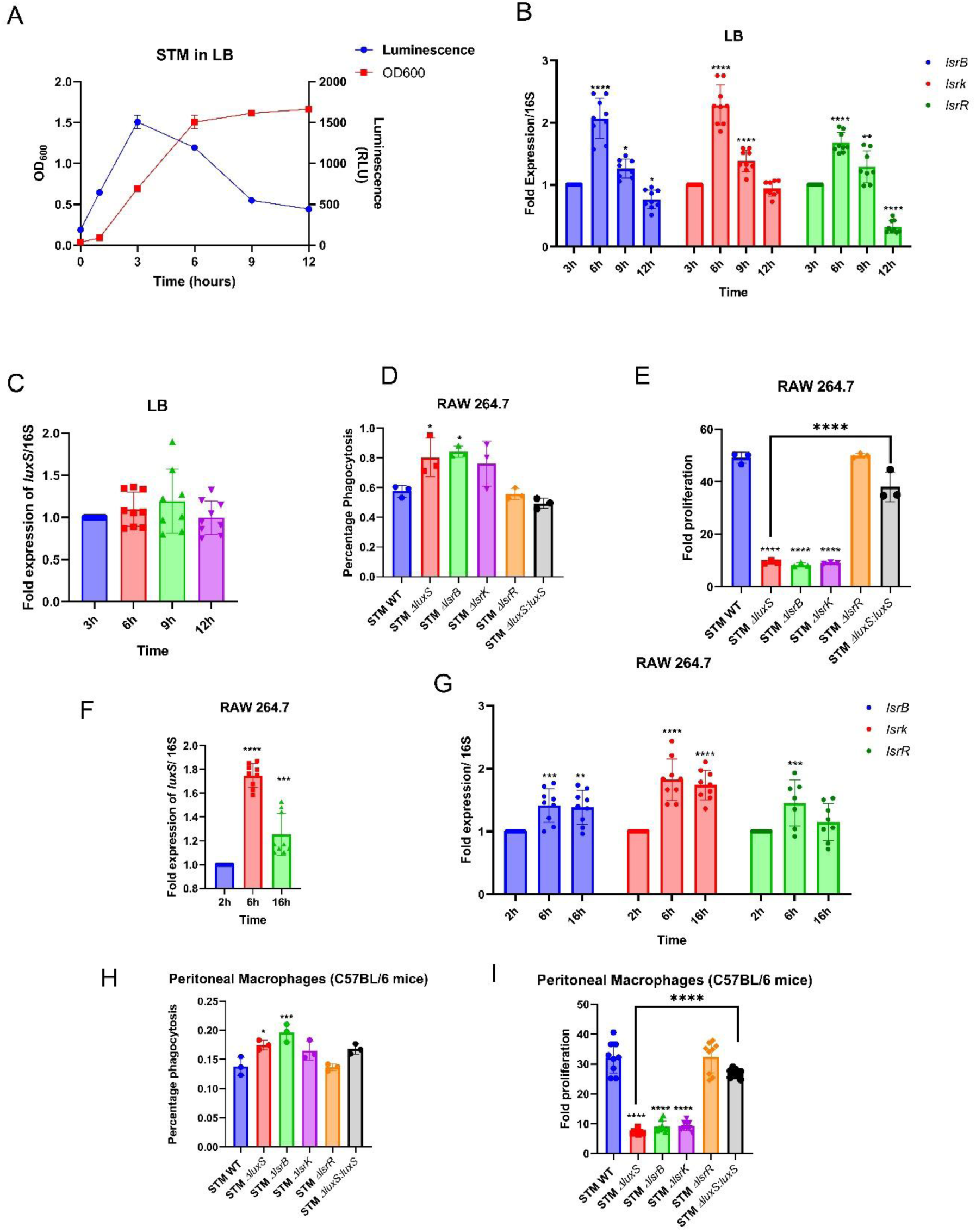
LuxS/AI-2 quorum sensing is required by *Salmonella* Typhimurium to survive in the macrophages. **(A)** *Salmonella* growth with AI-2 production in LB media. Represented as Mean*±*SD of N=3, n=4, **(B)** The mRNA expression of *lsr* operon genes *lsrB*, *lsrK*, and *lsrR* and **(C)** *luxS* gene expression in STM WT *in vitro* growth in LB. Represented as Mean*±*SD of N=3, n=3, **(D)** Percentage phagocytosis, and **(E)** Fold proliferation upon infection in RAW 264.7 macrophages. Represented as Mean*±*SEM of N=3, n=3, **(F)** *luxS* gene **(G)** *lsr* operon gene *lsrB*, *lsrk*, and *lsrR* expression in STM WT upon infection in RAW264.7 macrophages. Represented as Mean*±*SD of N=3, n=3 **(H)** Percentage phagocytosis and **(I)** Fold proliferation, upon infection in peritoneal macrophages. Represented as Mean*±*SEM of N=3, n=3. One-way ANOVA with Dunnet’s post-hoc test was used to analyze the data; p values **** p < 0.0001, *** p < 0.001, ** p<0.01, * p<0.05. Two-way Anova was used to analyze the grouped data; p values **** p < 0.0001, *** p < 0.001, ** p<0.01, * p<0.05

Further, we noted that the *luxS* and the *lsrB*, *lsrK,* and *lsrR* genes showed 1.5–2 fold upregulation post 6h of infection into RAW 264.7 macrophages till 16h **(Fig. 1F, G**). We further corroborated our results by using primary macrophages derived from the peritoneal lavage of C57BL/6J mice for infection. The *luxS, lsrB*, and *lsrK* mutants also exhibited an increased uptake by the primary macrophages, but their proliferation was compromised **(Fig. 1H, I)**. Furthermore, we checked for AI-2 production by STM WT, STMΔ*luxS*, and STMΔ*lsrB* upon their infection of RAW 264.7 cells and observed the AI-2 production by STM WT and STMΔ*lsrB* till 16h post-infection but not by STMΔ*luxS* **(Fig. S1D)**. Taken together, these findings imply that *Salmonella* Typhimurium requires LuxS/AI-2 signaling to survive in the hostile environment of macrophages.

### STM Δ*luxS* acquires AI-2 from STM WT during its *in vitro* growth but may not do so within its intravacuolar niche

To find out if STMΔ*luxS* can use AI-2 molecules produced by STM WT, we co-infected RAW 264.7 macrophages with STM WT and STMΔ*luxS* (1:1 ratio), and observed that STM WT outcompeted STMΔ*luxS,* hinting that STMΔ*luxS* may not be able to utilize the AI-2 from STM WT within host macrophages. **(Fig. 2A)**. However, when STM WT and STMΔ*luxS* were cocultured in LB media before infection into macrophages, STM WT and STMΔ*luxS* proliferated similarly in RAW 264.7 cells **(Fig. 2B)**. To check if STMΔ*luxS* can utilize AI-2 molecule produced by STM WT, we treated STMΔ*luxS* with the STM WT spent media and exogenous DPD molecule. Upon infection into RAW 264.7 macrophages, the treated STMΔ*luxS* proliferated similarly to STM WT, suggesting its ability to use the AI-2 from the spent media of STM WT or exogenous DPD **(Fig. 2C, D; Fig. S2A-D).** Next, we used a commercially available inhibitor (Z)-4-bromo-5-(bromomethyllene)-3-methylfran-2(5H) (BF-8) of LuxS/AI-2 signaling (18) (19) at different concentrations with STM WT and observed that while BF-8 did not affect STM WT growth *in vitro*, it attenuated its survival in RAW 264.7 macrophages similar to that observed with STMΔ*luxS* **(Fig. 2E, F; Fig. S2E)**. We also confirmed the light production and inhibition upon synthetic DPD and inhibitor BF-8 treatments by autoinducer assay **(Fig. S2F, G).** Infections into the peritoneal macrophages suggested that while exogenous DPD enhanced STMΔ*luxS* survival, BF-8 attenuated the STM WT proliferation **(Fig. S2H, I).** These findings imply that the *luxS* mutant can utilize exogenous AI-2 during its *in vitro* growth but not within its intravacuolar niche.

**Fig. 2:**
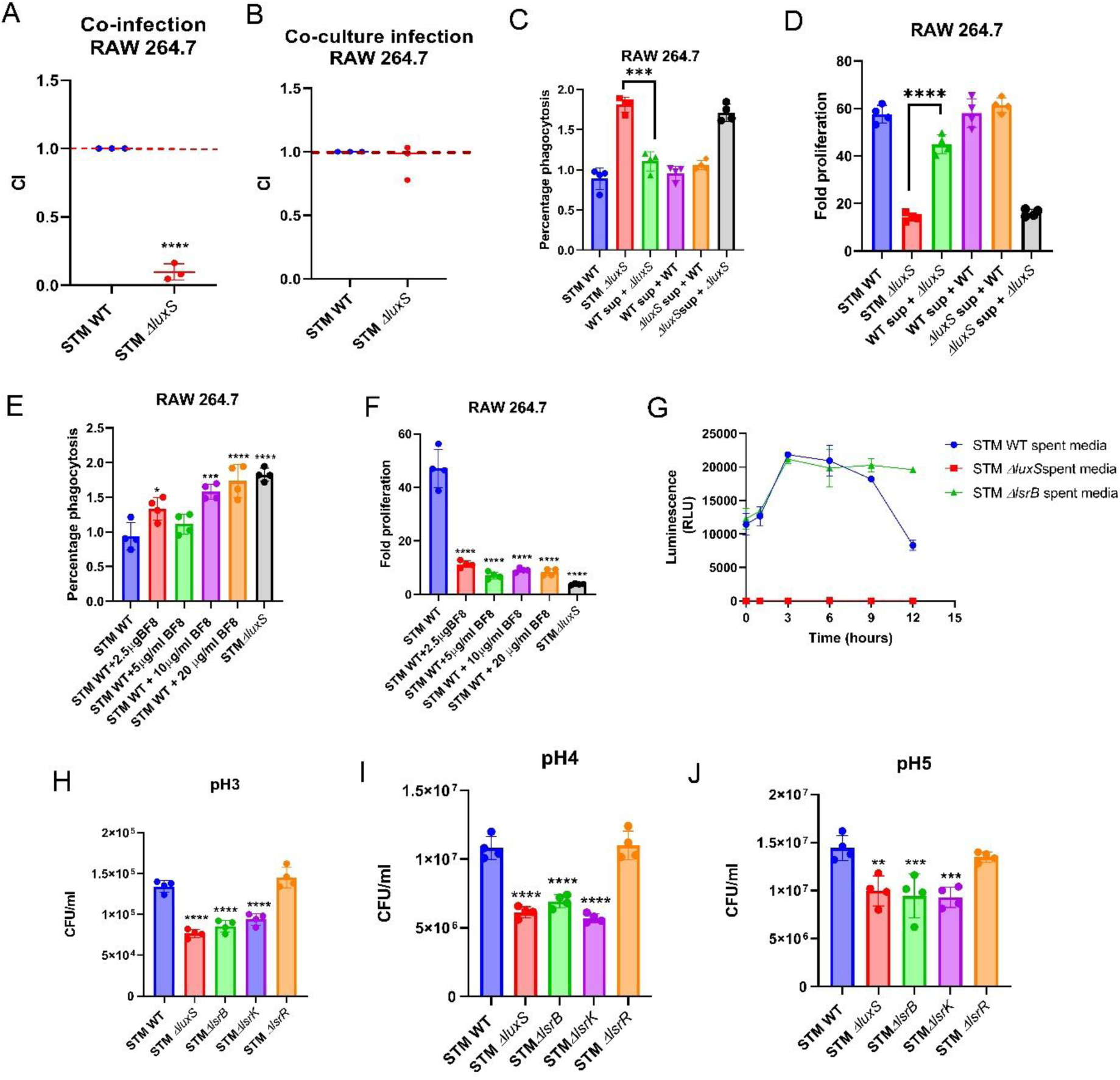
STM Δ*luxS* acquires AI-2 from STM WT during its *in vitro* growth but not do so within its intravacuolar niche. **(A)** Co-infection assay between STM WT and STMΔ*luxS upon* infection in RAW 264.7 macrophages at the ratio 1:1. Represented as Mean*±*SEM of N=3, n=3, **(B)** Co-culture grown STM WT and STMΔ*luxS* followed by infection in RAW 264.7 macrophages. Represented as Mean*±*SEM of N=3, n=3, **(C)** Percentage phagocytosis, **(D)** Fold proliferation of STM in RAW 264.7 macrophages upon 50% spent media treatment of STM WT and STMΔ*luxS*. Represented as Mean*±*SD of N=3, n=3, **(E)** Percentage phagocytosis, **(F)** Fold proliferation of STM in RAW 264.7 macrophages upon treatment of BF-8 inhibitor. Represented as Mean*±*SD of N=3, n=3. **(G)** AI-2 production by STM WT, STMΔ*luxS*, and STMΔ*lsrB* in F-media (pH 5). Represented as Mean*±*SD of N=3, n=3. STM WT, STM*ΔluxS*, STM*ΔlsrB*, STM*ΔlsrK,* and STM*ΔlsrR* survival in different ranges of pH(3-8) of PBS (H), pH 3 survival, **(I)** pH 4 survival, **(J)** pH 5 survival. Unpaired Student’s t-test was used to analyze the data; **** p < 0.0001, *** p < 0.001, ** p<0.01, * p<0.05. One-way ANOVA with Dunnet’s post-hoc test was used to analyze the data; p values **** p < 0.0001, *** p < 0.001, ** p<0.01, * p<0.05

### The loss of a functional AI-2 or its receptor-mediated sensing attenuates *Salmonella* survival at acidic pH

Quorum sensing pathways monitor changes in the density of the bacterial population and control the expression of genes involved in critical cellular processes, such as survival and pathogenicity (20). The mechanisms underlying bacterial quorum sensing and pH sensing have been thoroughly investigated and shown to be distinct/independent gene regulatory systems. Our study shows that environmental acidification is the critical physiological signal that upregulates AI-2 production by *Salmonella* **(Fig. 2G).** Hence, we hypothesize that LuxS/AI-2 signaling may coordinate with the pH sensing system of *Salmonella* to regulate survival in an acidic environment. Firstly, when we assessed the expression of mRNAs from *luxS* and *lsr* operon during its growth in F-media (mimics the acidic vacuolar condition), we observed ∼1.5-fold increases in *luxS* at the mid-log phase to the late-log phase **(Fig. S3A),** and *lsrB*, *lsrK,* and *lsrR* at the mid-log phase **(Fig. S3B)**. Moreover, we observed that STM stably produces AI-2 even during its growth in F-media (at pH 5), which is maximal during the mid to late-log phase **(Fig. 2G).** The luminescence production continued in STM*ΔlsrB* spent media even beyond 12 h (because there was no accumulation of AI-2 inside the STM *ΔlsrB*). However, there was no light production from the spent medium of STM*ΔluxS* **(Fig. 2G)**.

To underscore the role of AI-2 in facilitating survival in an acidic environment, we performed an acid survival assay and noted that STM WT, STM*ΔluxS*, ST *ΔlsrB*, STM*ΔlsrK*, and STM*ΔlsrR* survived equally well at pH6 to pH8 **(Fig. S3C)**. However, at pH3, pH4, and pH5, STM WT survived significantly better **(Fig. 2H-J; Fig. S3D-G)**. We conclude that LuxS/AI-2 signaling and communication facilitate *Salmonella’s* survival strategies in acidic environments under *in vitro* conditions.

### LuxS/AI-2 system tunes *phoP/phoQ* expression, thereby assisting in pH sensing and adaptation in *Salmonella*

Jesudhasan et al., 2010 determined that there is a differential gene expression between the STM WT and STMΔ*luxS* (21) pertaining to virulence, flagellar, and motility genes. A well-known two-component system(TCS) of PhoP/PhoQ is essential in mediating the acid tolerance response (ATR) in several Gram-negative bacteria, including *Salmonella* (22) (23). Thus, we conjectured that LuxS/AI-2 signaling modulates the *phoP/phoQ* gene expression, which, in turn, aids in the survival of *Salmonella* under an acidic environment. Thus, we questioned if phosphorylation of AI-2 is essential to regulate the expression of the downstream gene targets apart from the *lsr* operon **(Fig. 3A)**.

**Fig. 3:**
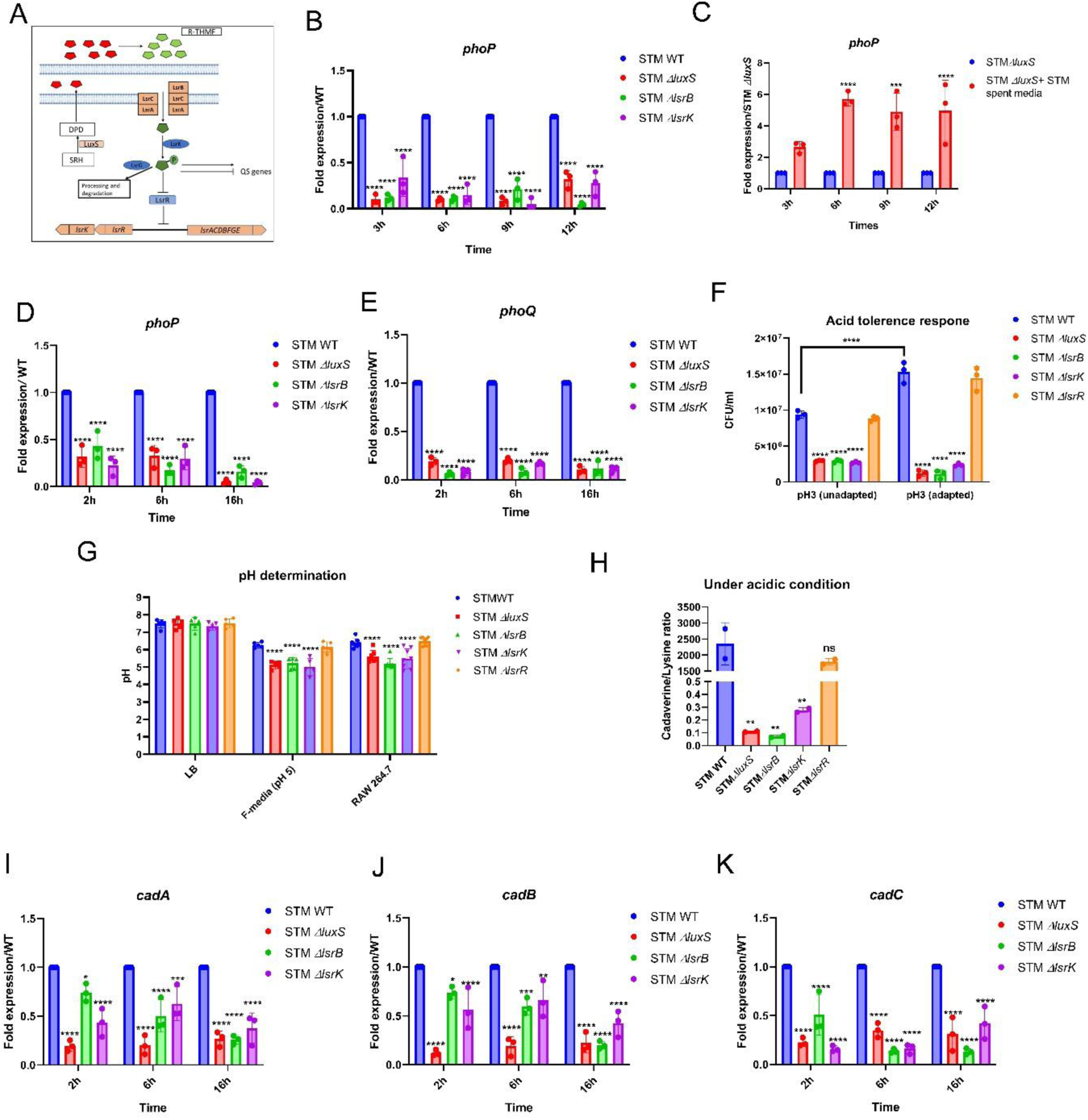
LuxS/AI-2 system tunes *phoP* expression, thereby assisting in pH sensing and adaptation in *Salmonella.* **(A)** Schematic of AI-2 synthesis and processing in *Salmonella* Typhimurium. **(B)** *phoP* gene expression in STM WT, STM*ΔluxS*, STM*ΔlsrB*, and STM*ΔlsrK in vitro* conditions LB media. Represented as Mean*±*SD of N=3, n=3. **(C)** mRNA expression of *phoP in* STM*ΔluxS* upon treatment of STM WT spent media. Represented as Mean*±*SD of N=3, n=3. **(D)** *phoP* **(E)** *phoQ* expression in STM WT, STM*ΔluxS*, STM*ΔlsrB*, and STM*ΔlsrK* upon infection in RAW 264.7 macrophages. Represented as Mean*±*SD of N=3, n=3. **(F)** STM WT, STM*ΔluxS*, STM*ΔlsrB*, STM*ΔlsrK*, and STM*ΔlsrR* acid adaptation and tolerance. Represented as Mean*±*SD of N=3, n=3. **(G)** Cytosolic pH determination of STM WT, STM*ΔluxS*, STM*ΔlsrB*, STM*ΔlsrK*, and STM*ΔlsrR* by phuji plasmid in LB Media, in F-media (pH 5) and upon infection RAW 264.7 macrophages. Represented as Mean*±*SD of N=3, n=4. **(H)** Ratio of cadaverine and lysine under acidic conditions (pH 5), N=2. **(I)** *cadA*, (**J)** *cadB,* and **(K)** *cadC*, mRNA expression upon infection in RAW 264.7 macrophages. Represented as Mean*±*SD of N=3, n=3. One-way ANOVA with Dunnet’s post-hoc test was used to analyze the data; p values **** p < 0.0001, *** p < 0.001, ** p<0.01, * p<0.05. Two-way ANOVA was used to analyze the grouped data; p values **** p < 0.0001, *** p < 0.001, ** p<0.01, * p<0.05

We observed that compared to STM WT, mRNA expression of *phoP* is downregulated in STM*ΔluxS*, STM*ΔlsrB*, and STM*ΔlsrK* in LB **(Fig. 3B; Fig. S4A)** and F-media **(S4B, D)**. Interestingly, the *phoP* expression was rescued when STM*ΔluxS* was grown in STM WT spent media **(Fig. 3C)**. Additionally, upon infection in macrophages, both *phoP* and *phoQ* were under-expressed in the mutants compared to STM WT **(Fig. 3D, E; Fig. S4C)**. Hence, we conclude that AI-2 sensing and the signaling cascade orchestrate the *phoP/phoQ* expression in *Salmonella* Typhimurium.

Next, we evaluated the ATR and noted that with a prior adaptation to pH 5, only STM WT and STM*ΔlsrR* were resilient and showed better survival upon exposure to pH 3 **(Fig. 3F)**. As the STM*ΔluxS*, STM*ΔlsrB*, and STM*ΔlsrK* exhibited a diminished ATR, we determined the intracellular pH of all the strains using a pH-sensitive pHuji plasmid. We observed that all the *Salmonella* strains maintained their cytosolic pH in the neutral LB media (pH 7). However, when grown in acidic F-media (pH 5), STM WT and STM*ΔlsrR* could maintain their cytosolic pH within the near-neutral range while cytosolic pH for STM*ΔluxS*, STM*ΔlsrB*, and STM*ΔlsrK* dropped to pH 5.1, 5.2, and 5.0, respectively and to 5.6, 5.4, and 5.2 (±0.2) respectively within the hostile macrophages **(Fig. 3G; Fig. S4E-I).**

The PhoP/PhoQ TCS aids in pH homeostasis via the *cadC/BA* system (transcriptional regulator, lysine/cadaverine antiporter and lysine decarboxylase), thereby promoting bacterial survival in mildly acidic environments (24, 25). The diminished mRNA expression of *cadC*, *cadB*, and *cadA* in STMΔ*phoP* upon infection in RAW 264.7 macrophages validated the above observation **(Fig. S5A-C)**. The lysine decarboxylase (CadA) quenches the H^+^ while converting lysine to cadaverine, which is exported to the extracellular milieu by CadB (24). Here, we noted the cadaverine/lysine ratio to be low in STM*ΔluxS*, STM*ΔlsrB*, and STM*ΔlsrK* compared to STM WT under an acidic pH of 5 **(Fig. 3H).** Additionally, in the mutants, the *cadA*, *cadB*, and *cadC* genes were under-expressed at 2h, 6h, and 16h post-infection into RAW 264.7 cells **(Fig. 3J, K, L).** From a mechanistic perspective, we conclude that LuxS/AI-2 signaling regulates ATR and maintains the cytosolic pH of *Salmonella* by regulating *cadC/AB* operon via *phoP/phoQ*.

### LuxS/AI-2 signaling regulates *phoP/phoQ* expression through LsrR interaction with the *phoP* promoter in *Salmonella*

We explored whether LsrR, the downstream regulator of the LuxS/AI-2 signaling pathway, could directly interact with the *phoP* promoter to influence its activity. The AlphaFold3 structure revealed that LsrR binds to both the DNA strands of the *lsr* promoter, but to only one of the strands of the *phoP* promoter (26). The interactions are mediated by residues Y25, T31, Q32, S33, R43, L44, K45, S47, and R48, all within a distance of <5 Å and a predicted aligned error (PAE) of <14 across multiple replicates **(Fig. 4A, B; Fig. S6A, B)**. These residues are located in the N-terminal domain, spanning positions 25-48, between alpha helices 1, 2, and 3. Notably, Y25, T31, and R43 contributed the most significant interactions, with R43 from helix 3 displaying the highest number of contact points (27). A covariance analysis of the multiple sequence alignment revealed that R43 most frequently co-occurs with Y25, an interacting residue located in helix 1, with the highest probability among all analyzed residues (28). Given the strong interactions of R43 and Y25 with the *lsr* and *phoP* promoters, we generated Y25A, R43A, and Y25A/R43A mutants and used AlphaFold3 to predict structural impacts. The AlphaFold structures indicated that the mutations severely disrupted promoter interactions, with PAE values exceeding 18 and pLDDT scores for all the DNA-contacting residues dropping below 30. Additionally, we performed NPT ensemble molecular dynamics simulations for the mutant protein-DNA complex, which revealed a significant weakening of interactions. We validated our findings *in vitro* through electrophoretic mobility shift assays (EMSA) using purified wild-type LsrR and its Y25A, R43A, and Y25A/R43A mutants to assess their binding to HPLC-purified *lsr* and *phoP* promoters **(Fig. 4D, E; Fig. S6C-F)**. WT LsrR bound both promoters, Y25A showed reduced binding, R43A further reduced it, and the double mutant lost nearly all binding at 200mM NaCl **(Fig. S7A, B)**. A similar result was observed for the interaction of LsrR with a single-stranded promoter sequence **(Fig. S7C-F),** and a random DNA sequence did not show any interaction (**Fig. S7G)**. To assess regulatory effects during infection, we generated a *luxS/lsrR* double knockout and complemented it with *lsrR* (WT) and mutant *lsrR*. While the double knockout strain proliferated similarly to WT STM, *lsrR* complementation reduced proliferation. The Y25A/R43A mutant restored survival to knockout levels **(Fig. 4E, F)**, indicating LsrR’s binding to *lsr* and *phoP* promoters acts as a negative regulator of *Salmonella* pathogenicity.

**Fig. 4:**
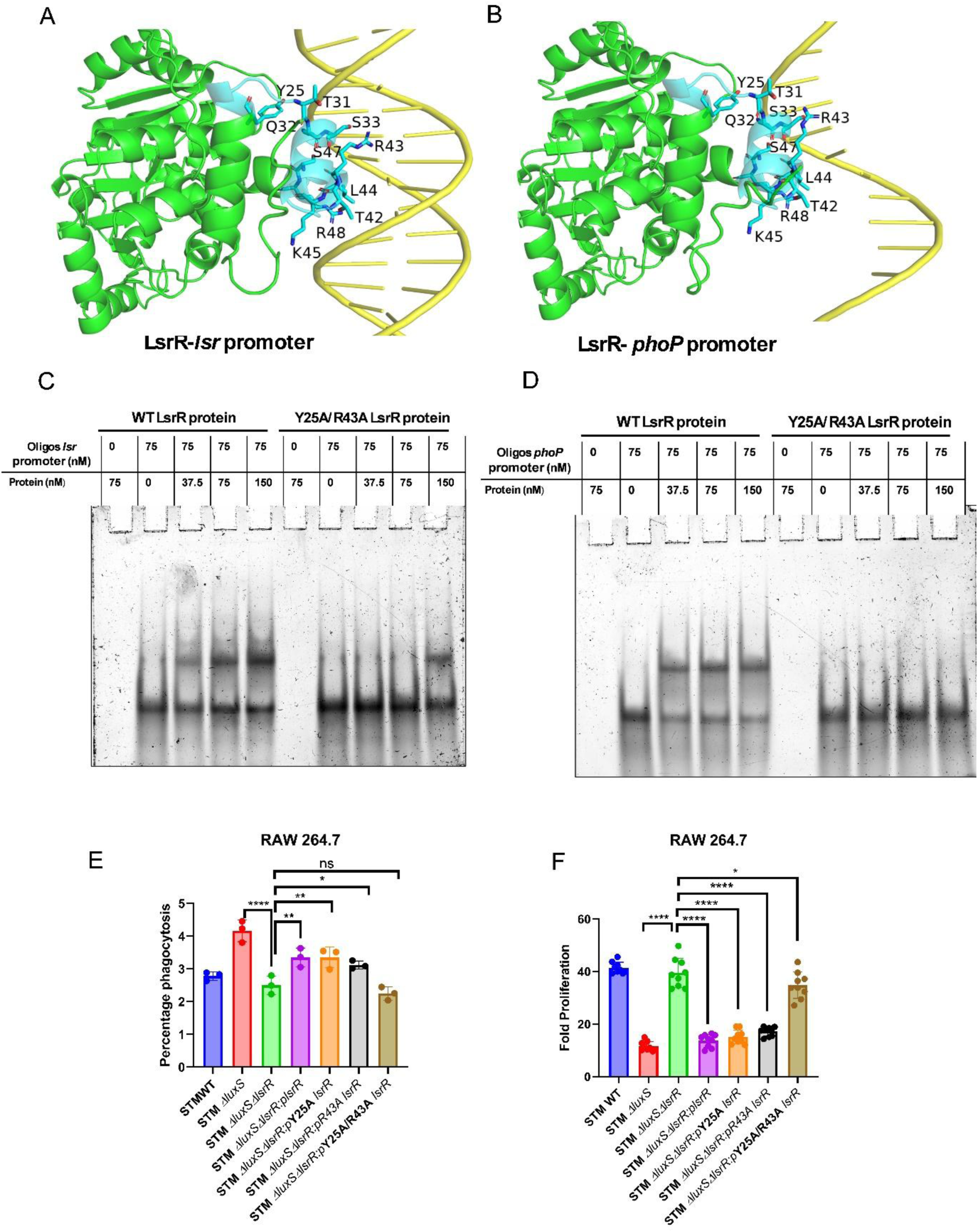
LuxS/AI-2 signaling regulates *phoP/phoQ* expression through LsrR interaction with the *phoP* promoter in *Salmonella.* AlphaFold3 structural models visualised using pymol depicting LsrR (green) interactions with the *phoP* and *lsr* promoters (yellow). The N-terminal domain residues, spanning helices 1, 2, and 3, are highlighted in blue, and the key interacting residues of LsrR are represented with a ball and stick model **(A, B)**. Electrophoretic mobility shift assays (EMSA) assessing DNA binding of wild-type LsrR and its Y25A/R43A mutant to *lsr* and *phoP* promoters (**C, D)**. Intracellular survival assay (ICSA) with mutated Y25A, R43A, and Y25A/R43A *lsrR* cloned in pQE-60 complemented strains **(E)** phagocytosis and **(F)** Fold proliferation. Represented as Mean*±*SD of N=3, n=3. One-way ANOVA with Dunnet’s post-hoc test was used to analyze the data; p values **** p < 0.0001, *** p < 0.001, ** p<0.01, * p<0.05.

### AI-2 orchestrates the *Salmonella* Pathogenicity Island-2 gene expression by sensing low pH

PhoP has been shown to control the TCS SsrA/SsrB at the transcription level, which are the master regulators of the SPI-2 genes (29). The expression of *ssrA* and *ssrB* genes was also downregulated in STMΔ*phoP* upon infection into RAW 264.7 cells **(Fig. S8A, B)**. Thus, we hypothesized that LuxS/AI-2 signaling might control the SsrB/SsrA system via PhoP/PhoQ. Upon infection into RAW 264.7 macrophages, the mRNA expression of *ssrB* was downregulated from early stage post-infection to late stage post-infection in macrophages **(Fig. 5A)**. On the other hand, *ssrA* was downregulated at a late stage post-infection **(Fig. 5B)**. Furthermore, we observed expression of *ssaV* a Type 3 secretion system (T3SS) needle complex protein, encoded by SPI-2 and *spiC* an SPI-2 encoded effector molecule were under-expressed in the absence of *luxS, lsrB,* and *lsrK* genes in STM upon infection into RAW 264.7 macrophages **(Fig. 5C, D).**

**Fig. 5:**
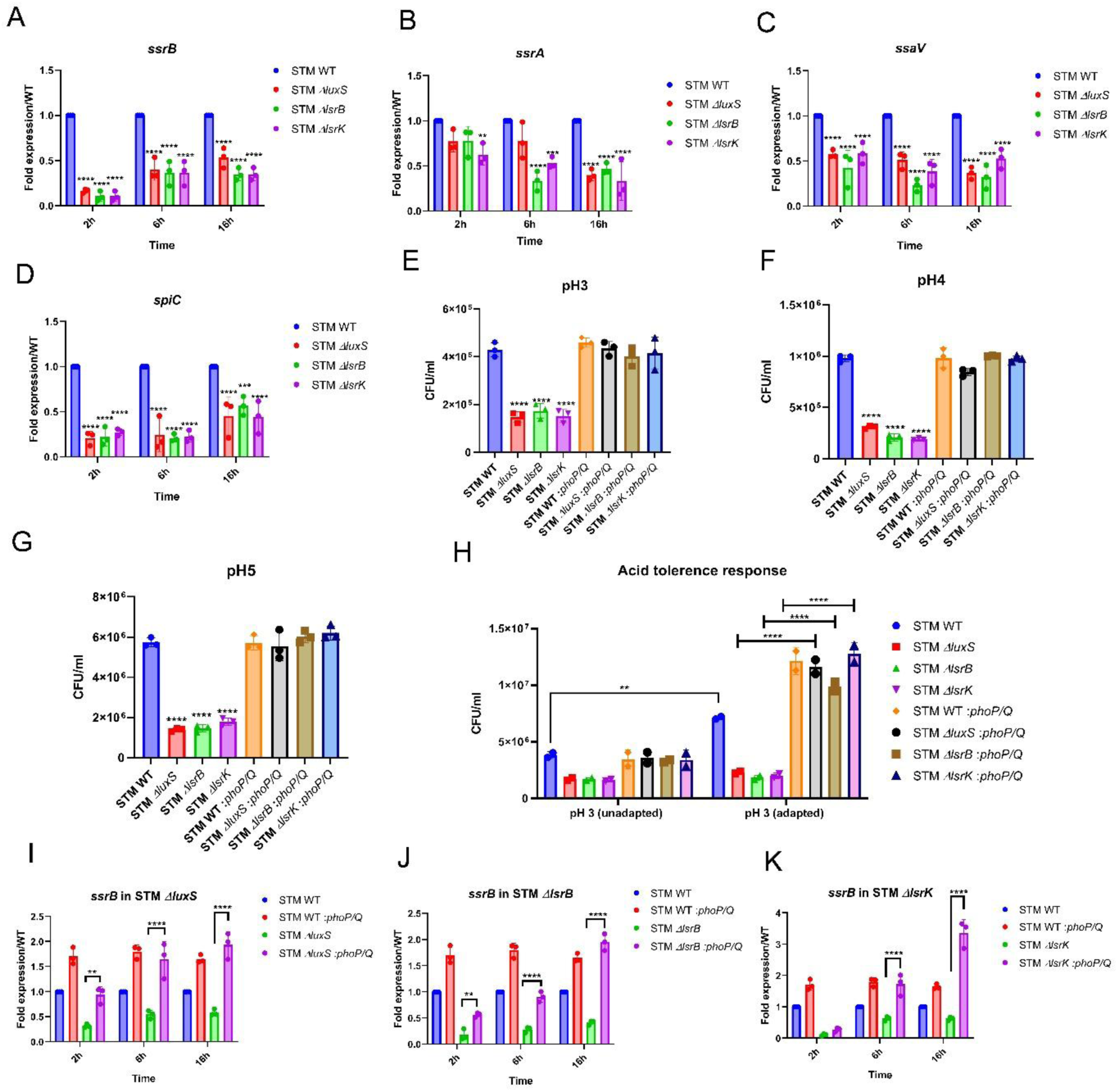
AI-2 orchestrates the *Salmonella* pathogenicity island-2 gene expression by sensing low pH. **(A)***ssrB* **(B)** *ssrA* **(C)** *ssaV* **(D)** *spiC,* mRNA expression in STM WT, STM*ΔluxS*, STM*ΔlsrB*, and STM*ΔlsrK* upon infection in RAW 264.7 macrophages. Represented as Mean*±*SD of N=3, n=3. STM WT, STM*ΔluxS*, STM*ΔlsrB*, STM*ΔlsrK,* STM WT: *phoP*, STM*ΔluxS*: *phoP*, STM*ΔlsrB*: *phoP*, STM*ΔlsrK*: *phoP* survival in PBS at **(E)** pH3, **(F)** pH4, **(G)** pH5. Represented as Mean*±*SD of N=3, n=2. **(H)** STM WT, STM*ΔluxS*, STM*ΔlsrB*, STM*ΔlsrK,* STM WT:*phoP*, STM*ΔluxS*:*phoP*, STMΔlsrB:*phoP*, STM*ΔlsrK*: *phoP* acid adaptation and tolerance. Represented as Mean*±*SD of N=3, n=2. *ssrB* mRNA expression in *phoP*/*phoQ* cloned strain of **(I)** STM*ΔluxS,* **(J)** STM*ΔlsrB* and **(K)** STM*ΔlsrK.* Represented as Mean*±*SD of N=3, n=3. One-way ANOVA with Dunnet’s post-hoc test was used to analyze the data; p values **** p < 0.0001, *** p < 0.001, ** p<0.01, * p<0.05. Two-way ANOVA was used to analyze the grouped data; p values **** p < 0.0001, *** p < 0.001, ** p<0.01, * p<0.05

So far, our findings convey that the LuxS/AI-2 signaling regulates *phoP/phoQ* genes and, thereby, the SPI-2 cluster in *Salmonella*. However, whether AI-2 directly or the AI-2-PhoP/PhoQ axis controls the SPI-2 genes remains unclear. Thus, we cloned the *phoP*/*phoQ* operon under a non-native promoter in the pQE60. We confirmed the *phoP* mRNA overexpression level in STM WT:pQE60-*phoP*/*phoQ* and STM*ΔluxS:*pQE60-*phoP*/*phoQ* upon infection in RAW 264.7 **(Fig. S8C, D)**. Using these complemented *Salmonella* strains, we observed that STM*ΔluxS*, STM*ΔlsrB*, and STM*ΔlsrK* with a *phoP*/*phoQ* cloned under a non-native promoter survived equally at acidic pH3, 4, and 5 as STM WT **(Fig. 5E-G)**. Furthermore, the STM*ΔluxS*, STM*ΔlsrB*, and STM*ΔlsrK* with cloned *phoP*/*phoQ* showed enhanced ATR response **(Fig. 5H)**. Thus, when *phoP/phoQ* is not under its native promoter, the regulation by LuxS/AI-2 is not required. Also, *ssrB* and *ssrA* expression are significantly higher in *phoP*/*phoQ* complemented knockout strains compared to STM*ΔluxS*, STM*ΔlsrB*, and STM*ΔlsrK* strains **(Fig. 5I-K; Fig. S8E-G)**. Cumulatively, these results suggest that, indeed, the LuxS/AI-2 signalling controls ATR by regulating the *phoP/phoQ* genes.

### The absence of AI-2 mediated signaling compromises the *in vivo* colonization of *Salmonella* in mice

The gastrointestinal tract is a diverse and dynamic environment in which bacterial species communicate with each other to coordinate the expression of various genes (30). While many quorum sensing signals are species-specific, AI-2 signalling is known as universal quorum sensing signaling. Because AI-2 produced by one species can affect the expression of genes in another, this signal can promote communication across species and allow bacteria to alter behaviors like pathogenicity, and the production of biofilms among various species (31). Because of this characteristic, AI-2 is an excellent choice to mediate interactions between cells in the mammalian gut, where hundreds of bacterial species live and interact. It has already been determined that several gut-associated bacteria encode LuxS (the AI-2 synthase) or generate AI-2 (32). Of the major families dominating the gut consortium, more than 80% of Firmicutes have been detected to encode the LuxS enzyme (33). While members of the families *Barnesiellaceae* and *Muribaculaceae* are predominant producers of AI-2 in the gut (34). Thus, we aimed to understand the role of LuxS/AI-2 signalling in regulating *Salmonella* pathogenesis in *in vivo* mice models.

We infected C57BL/6J mice by orally gavaging with STM WT and knockout strains at a CFU of 10^7^ per mouse **(Fig. 6A**). We observed that STM*ΔluxS*, STM*ΔlsrB*, and STM*ΔlsrK* showed a significantly lower organ burden in the intestine, MLN, spleen, and liver and lower bacteraemia upon oral gavage **(Fig. 6B–F)**. Oral gavage mimics the physiological route of *Salmonella* infection into its host, and *Salmonella* is required to be able to breach the intestinal barrier successfully. The lower organ colonization in STM*ΔluxS*, STM*ΔlsrB*, and STM*ΔlsrK* can be explained by their inability to cross the gut epithelial barrier. Next, we bypassed the gut epithelial barrier by infecting C57BL/6 mice intraperitoneally and found that STM*ΔluxS,* STM*ΔlsrB, and* STM*ΔlsrK* still exhibited reduced colonization in the spleen and liver and less dissemination in blood than STM WT **(Fig. S9A-D).** Furthermore, upon infection of mice by oral gavaging at a CFU of 10^8^ per mouse, we noted that the mice infected with STM WT and STM*ΔlsrR* succumbed to death as early as the 6th day of post-infection, while STM*ΔluxS*, STM*ΔlsrB*, and STM*ΔlsrK* infected mice survived longer **(Fig. 6G-H)** with less weight reduction **(Fig. S9E)**. Further, we observed that the BF-8 inhibitor at 4 mg/kg reduced the STM WT colonization to different sites of infection in C57BL/6 mice **(Fig. 6I-N**) and improved the survival of mice with delayed weight reduction compared to untreated mice **(Fig. 6O, P; Fig. S9F).** Liver tissue histopathology results also suggest that treatment with BF-8 reduced the disease score **(Fig. S9G)**. Our results show that LuxS/AI-2 signaling is critical for the *in vivo* pathogenesis and virulence of *Salmonella* Typhimurium.

**Fig. 6:**
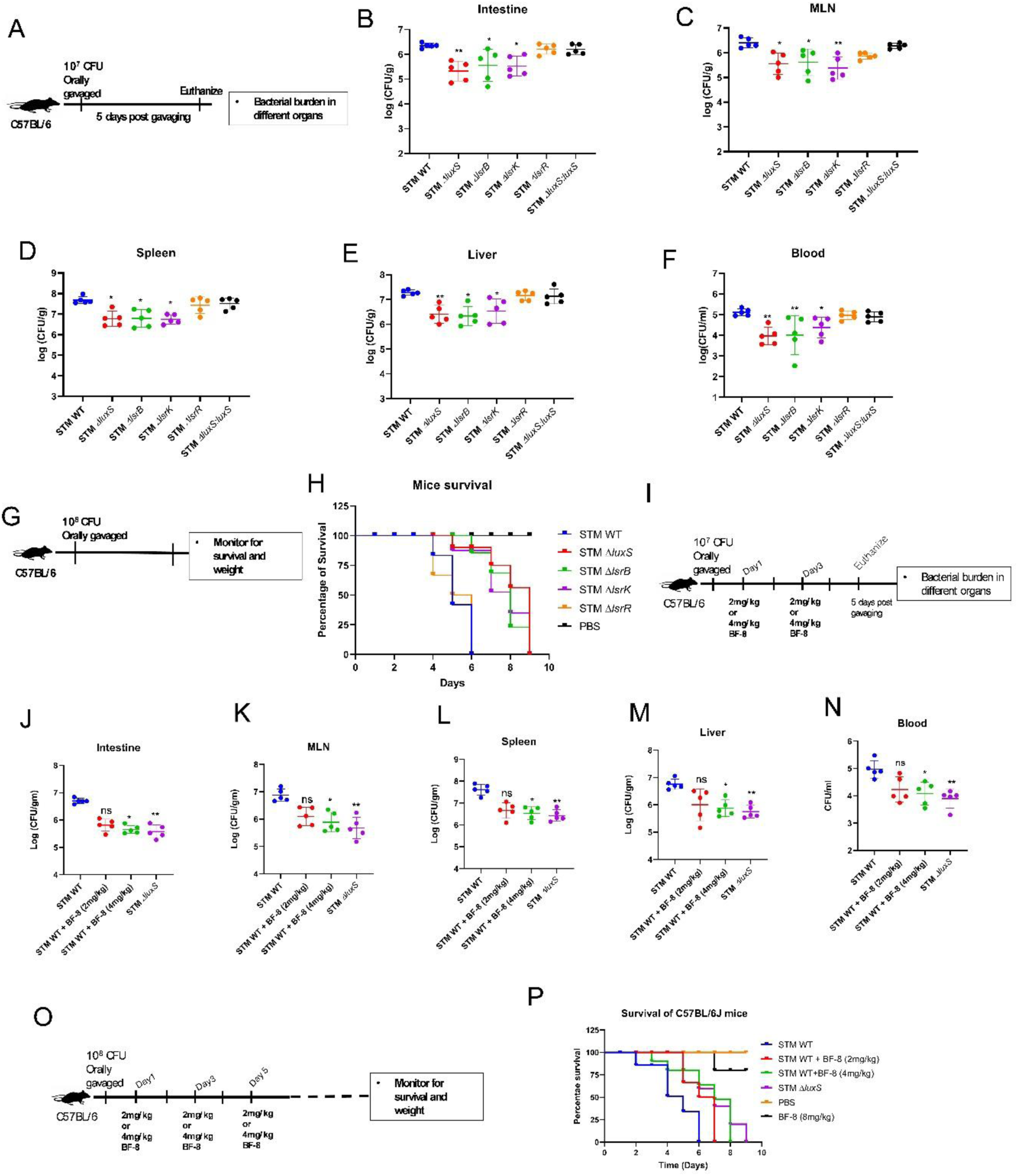
The absence of AI-2 mediated signaling compromises the *in vivo* colonization of *Salmonella.* **(A)** The experimental protocol for organ burden in C57BL/6J mice by orally gavaging 10^7^ CFU per mouse. The organ burden post 5 days of oral gavage in **(B)** Intestine, **(C)**Mesenteric lymph node (MLN), **(D)** Spleen, **(E)** Liver, and dissemination in **(F)** Blood. Represented as Mean*±*SD of N=2, n=5 **(G)** The experimental protocol for C57BL/6J mice survival by orally gavaging 10^8^ CFU of STM per mouse. **(H)** Percentage survival of Mice upon infection with STM WT, STM*ΔluxS*, STM*ΔlsrB*, STM*ΔlsrK,* STM*ΔlsrR* and STM*ΔluxS:luxS.* (**I)** The experimental protocol for organ burden in C57BL/6J mice by orally gavaging 10^7^ CFU per mouse upon treatment of BF-8 inhibitor (2mg/kg and 4mg/kg through the Intraperitoneal route). The organ burden post 5 days of oral gavage in **(J)** Intestine, **(K)**Mesenteric lymph node (MLN), **(L)** Spleen, **(M)** Liver, and blood dissemination in **(N)** Blood. Represented as Mean*±*SD of N=2, n=5. **(O)** The experimental protocol of mice survival in C57BL/6J mice by orally gavaging 10^8^ CFU per mouse upon treatment of BF-8 inhibitor (2mg/kg and 4mg/kg through the Intraperitoneal route). **(P)** Percentage survival of Mice upon infection with STM WT, STM WT upon treatment, and STM*ΔluxS.* Non- parametric One-way ANOVA (Kruskal Wallis) with Dunn’s posthoc test was used to analyze organ burden in mice.

## Discussion

The human body serves as a host for a vast array of microorganisms collectively known as the normal microbiota. These microbes coexist closely with their host and often offer significant health benefits, especially within the intestine. These microbes supplement our metabolism by producing essential vitamins like B and K, generating vital compounds such as short-chain fatty acids (SCFAs). These microbial functions play a key role in maintaining intestinal homeostasis, protecting and defending against both gut and systemic infections(35, 36).

Understanding how these microbial interactions influence health is critical, especially given the rising prevalence of infectious and chronic diseases linked to disruptions in the microbiota, also known as dysbiosis. One major trigger of dysbiosis is the use of broad-spectrum antibiotics and the use of secondary metabolites or chemical signals by pathogens, which can drastically alter the composition and diversity of gut bacteria(37, 38). These changes often reduce beneficial taxa while allowing harmful ones to proliferate. This illustrates the principle of colonization of a pathogen to establish an infection. This is true in the case of the enteric pathogen *Salmonella.* In the Oligo-mouse-microbiota (OMM^12^) model of infection, *Salmonella* induces compositional changes in the synthetic consortium (39). A previous study showed that the OMM^12^ mice model provides less colonisation resistance to *Salmonella* infection (40). It can be perceived that the ability of this pathogen to cause microbial shifts via distinct mechanisms is a stealthy trategy to evade colonisation resistance in the gut.

Beyond metabolism, behaviors such as motility, surface adherence, and biofilm formation further shape microbial dynamics. These processes are often coordinated through quorum sensing a communication system where bacteria release chemical signals called autoinducers to monitor population density and regulate gene expression collectively(20). Given the multispecies nature of this environment, cross-species signaling likely enables microbes to fine-tune their behaviors in response to social and environmental cues. Studies show that AI-2 facilitates interspecies communication and modulates behavior across species lines, leading to the hypothesis that AI-2 plays a central role in regulating microbial interactions and community dynamics in the gut(41).

The aspects of the bacterial lifestyle and pathogenesis rely on the bacterial quorum and community attributes, like for *Salmonella*, the infectious dose to cause the illness in humans is more than 10,000 CFU(42). Autoinducers are chemical signal molecules that underpin bacterial communication and differ amongst species, just like the different languages that exist across the human civilization. Here, we report that as a successful enteric pathogen, *Salmonella* can use the AI-2 molecule to coordinate and break the colonization resistance in the gut. AI-2 synthesis and *lsr* operon are concurrently regulated under neutral and acidic conditions in *Salmonella,* aiding in its survival. The phosphorylation of AI-2 by LsrK is crucial for its function (43) and is validated by our study, where we observe that deletion of *lsrK* renders *Salmonella* incapable of surviving in macrophages and incapable of upregulating *phoP* expression.

The gastrointestinal tract presents with a dynamic environment where microbes compete for space and nutrients through various mechanisms. Many gut-associated bacteria encode LuxS or produce AI-2, including a significant proportion of Firmicutes and Proteobacteria, and some species of Bacteroidetes and Actinobacteria (38). *Salmonella* might benefit from the AI-2 produced by the gut commensals in the intestinal lumen. Our *in vivo* studies justify it to some extent with compromised colonization of the primary and secondary sites of infection upon abrogation of LuxS/AI-2 signaling in *Salmonella*. Although the neighboring bacteria might produce AI-2 in the intestinal niche, STMΔ*lsrB* and STMΔ*lsrK* cannot utilize AI-2 to mediate the downstream signaling cascades. Thus, we underscore the critical function of individual members of this cascade, the AI-2 synthesizing enzyme, receptor, and kinase enzyme. Our study of baceterial strain co-culture prior to infection further suggests the ability of *Salmonella* to utilise the microbiota produced AI-2 molecules in the gut.

During intestinal colonization, a prime strategy of this pathogen is to activate the ATR systems to endure the harsh environment. The four major regulators that control relevant stress responses in *Salmonella* are RpoS, PhoPQ, Fur, and OmpR/EnvZ (44). We describe a mechanism that AI-2 signaling tunes the expression of the two-component system *phoP/Q*, which helps in ATR. Relating to intravacuolar survival, studies show that the acidic pH of the vacuole is necessary for the assembly of T3SS encoded by SPI-2, but not for the effector molecule’s secretion (45). Yet, it is essential to maintain the neutral pH of the bacterial cytoplasm to secrete the effector molecule, which we found to be lost upon abrogation of AI-2 signaling (46). Thus, *Salmonella* regulates its cytosolic pH in acidic environments via the AI-2 signaling and tries to sustain its intracellular pH near neutral to maintain the regular cellular functions and secretion of SPI-2 effectors for its survival and proliferation within the macrophages. Moreover, *Salmonella* AI-2 phosphorylation by LsrK is a key process in controlling the downstream targets. While our study elucidates the mechanism of acid tolerance and macrophage survival via AI-2 signaling in *Salmonella*, it has some limitations. We cannot conclusively determine whether STMΔ*luxS* utilizes AI-2 from STM WT in the coinfection study in RAW 264.7 cells **(Fig. 2A)**. Furthermore, the study does not confirm whether both bacterial strains infect the same cell, as observations reflect a global phenotype. However, it might be possible that STM *ΔluxS* cannot utilize AI-2 even when infecting the same cell due to a repressed *lsr* operon.

We broaden the horizon of LsrR regulation of gene expression beyond the *lsr* operon by unraveling the binding of LsrR to the *phoP* promoter via its Y25 and R43. Interestingly, these insights also shed light on an unusual interaction of the LsrR with a single strand of the DNA at the promoter site. This not only identifies the mechanism but also paves the way to a “secret garden” of genes that may be regulated by LsrR. While this observation is in *Salmonella,* it can be extrapolated to other bacteria that utilise a similar AI-2 signalling cascade. Overall, we identify a regulatory network via AI-2 and PhoP/PhoQ TCS, leading to a remarkable survival strategy for *Salmonella* under acidic pH. A proposed strategy for controlling *Salmonella* infections focuses not on bactericidal activity, as in the case of conventional antimicrobials, but rather on disrupting a key functional process critical to the pathogen’s infectious lifecycle. One promising target is the communication, wherein *Salmonella* uses it to actively survive in the acidic environment of the stomach, colonizes in the intestine, followed by a secondary site of infection, and survives in the hostile niche of macrophages, which is an essential step for successful pathogenesis. These virulence process depends on the coordinated regulation of numerous virulence genes, which are modulated through an intricate network of genetic and environmental signals. Our *in-vivo* study suggests that inhibition of LuxS/AI-2 signaling attenuates *Salmonella* colonization. Our findings highlight and open promising avenues to target the bacterial communication and AI-2 signaling, for designing antibacterial strategies to limit *Salmonella* infection.

## Ethics statement

All experiments comply with the rules set forth by the Indian Institute of Science, Bangalore’s IEAC. The approved protocol number is CAF/ Ethics/852/2021. The Committee for Control and Supervision of Experiments on Animals (CPCSEA), a statutory committee established under Chapter 4, Section 15 (1) of the Prevention of Cruelty to Animals Act 1960, and National Animal Care provided guidelines that were meticulously adhered to during all animal experiments, all of which were approved by the Institutional Animal Ethics Committee. (Registration No. 435 48/1999/ CPCSEA). The Institutional Animal Ethics Committee approved every animal experiment, and the National Animal Care Guidelines were meticulously followed.

## Author contributions

AS and DC conceived the study. AS and DC designed experiments. AS and AVN performed experiments. SA conducted structural analysis, and AS and SA performed interaction studies under the supervision of UV. SM supervised the synthetic DPD molecule synthesis. SD and SK synthesized the synthetic DPD molecule. RSR analyzed the tissue histopathology. AS analyzed the data, prepared the figures, and wrote the manuscript draft. AS, AVN, SA, SM, UV, and DC reviewed and edited the manuscript. DC supervised the work. All the authors read and approved the manuscript.

## Acknowledgment

Dr. Shashank Tripathi (CIDR, IISc Bangalore) and Dr. Sandeep M Eswarappa (Dept. of Biochemistry, IISc Bangalore) are duly acknowledged for the Luminometer. Divisional Mass Spectrometry facility IISc and Mrs. Sunita Joshi for the MS analysis. The Departmental Confocal Facility, Departmental Real-Time PCR Facility, Divisional Flowcytometry Facility, and Central Animal Facility at IISc are duly acknowledged. Mr Sumith and Ms Navya are acknowledged for their help in image acquisition. Dr. Ritika Chatterjee and Ms. Sagrika are acknowledged for technical help. Schematic images were created with BioRender (BioRender.com).

## Data availability statement

The data that support the findings of this study are available from the corresponding author (DC) upon request.

