## Supplementary material for "The quorum-sensing lexicon of *Salmonella* ameliorates acid stress in the host by a non-canonical mechanism": Suplementary Information

### Supplementary Text

### Chemical synthesis of acetonide-protected (*S*)-4,5-dihydroxy-2,3-petanedione (DPD)

#### A. General information:

Infrared (FT-IR) spectra were recorded on a Bruker Alfa FT-IR, $v_{max}$ in $cm^{-1}$ and the bands are characterized as broad (br), strong (s), medium (m), and weak (w). NMR spectra were recorded on Bruker Ultrashield spectrometer at 400 MHz (for $^{1}$H-NMR) and 100 MHz (for $^{13}$C-NMR). Chemical shifts are reported in ppm from tetramethylsilane with the solvent resonance as internal standard [$CDCl_3$: δ 7.26, $CD_3OD$: δ 3.31, $(CD_3)_2SO$: δ 2.50 for $^{1}$H-NMR and $CDCl_3$: δ 77.16, $CD_3OD$: δ 49.00, $(CD_3)_2SO$: δ 39.52 for $^{13}$C-NMR]. For $^{1}$H-NMR, data are reported as follows: chemical shift, multiplicity (s = singlet, d = doublet, dd = double doublet, ddd = doublet of doublet of doublets, t = triplet, q = quartet, sep = septet, br = broad, m = multiplet), coupling constants (Hz) and integration. High resolution mass spectrometry was performed on Waters XEVO G2-XS QTof instrument. Optical rotations were measured on a JASCO P-2000 polarimeter. Melting points were measured in open glass capillary using Büchi M-560 melting point apparatus. Enantiomeric ratios were determined by Shimadzu LC-20AD HPLC instrument and SPD-20A Diode Array Detector using stationary phase chiral columns (25 cm × 0.46 cm) in comparison with authentic racemic compounds.

#### B. Procedure for the preparation of (*S*)-2,3-dihydroxypropanoate 2

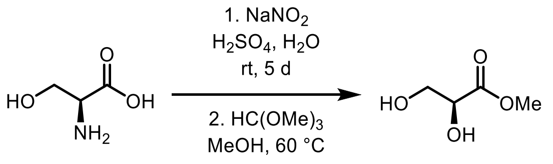


*L*-serine, **1**      **2**

(*S*)-2,3-Dihydroxypropanoate **2** was prepared using modified literature procedure(1). To an ice-cooled solution of *L*-serine **1** (15 g, 142.7 mmol) in 30 ml $H_2O$, $H_2SO_4$ (5M, 45 mL) was added, followed by slow addition of $NaNO_2$ (15.3 g in 27.0 mL water). The reaction mixture was then stirred at ambient temperature for 5 h. Another portion of aqueous $NaNO_2$ (6M, 27 mL, 12.4 g in 27.0 mL water) was added at 0 °C. The reaction mixture was stirred at ambient temperature for 3 d. The reaction mixture was cooled to 0 °C and $H_2SO_4$ (5M, 45mL) was added, followed by $NaNO_2$ (15.3 g in 27.0 mL water) slowly and stirred at ambient temperature for 2 d. The reaction mixture was concentrated under reduced pressure (70°C, 100-200 mbar). The resulting solution was treated with an aqueous NaOH solution (10 M) until neutralization. A mixture of MeOH/acetone (3:1, 400 mL) was added and passed through a pad of celite and repeated the process 7-10 times. Toluene was added (20 mL) for toluene-water azeotrope. The residue was dissolved in 60 mL of MeOH and 15-20 mL of trimethyl orthoformate (15.3g, 221.7 mmol, 1.55 equiv) was added. After that conc. $H_2SO_4$ was added until pH 1. The mixture was stirred at 60 °C for 1 h. Reaction was quenched by NaOMe at 0 °C. After filtration, solvent was evaporated, and the residue was purified by silica-gel column chromatography (10% Methanol-ethyl acetate) to obtain the (*S*)-2,3-dihydroxypropanoate **2** as a colorless liquid (10.1 g, 84.10 mmol, 59% yield); **$^1$H-NMR (400 MHz, $CDCl_3$):** δ 4.24 (t, *J* = 4.2,3.3 Hz, 1H), 4.1 (s, 2H), 3.81 (d, *J* = 3.1 Hz, 1H), 3.77 (d, *J* = 4.4 Hz, 1H), 3.72 (s, 3H); **$^{13}$C-NMR (100 MHz, $CDCl_3$):** δ 173.6, 71.8, 64.1, 52.9.

**C. Procedure for acetonide protection of (*S*)-2,3-dihydroxypropanoate 2**

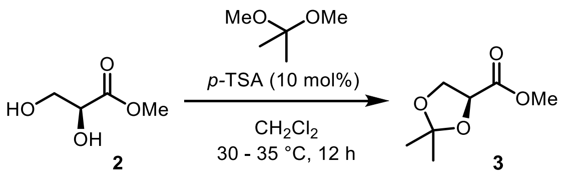

The acetonide protected methyl ester **3** was prepared using a literature procedure(2). In an oven dried 250 mL round bottom flask, **2** (8.7g, 72.75 mmol) was taken in 40 mL of dry $CH_2Cl_2$ and 2,2-dimethoxypropane (18mL, 145.5 mmol, 2.0 equiv) was added. The resulting solution was stirred at 0 °C for 5 min, following which 1.4g (7.2 mmol, 0.1 equiv) of *p*-TSA was added and stirring was continued at 0 °C for 5 min. The reaction mixture was then stirred at 30-35 °C for 12 h. The solvent was evaporated, and the residue was purified by vacuum distillation using 15 cm long Vigreux column (oil bath temperature 130-140 °C, 14 mbar pressure) to get acetonide **3** as a yellow liquid (8.3 g, 51.8 mmol, 71% yield); **$^1$H-NMR (400 MHz, $CDCl_3$):** δ 4.53 (dd, *J* = 7.1, 5.4 Hz, 1H), 4.17 (dd, *J* = 8.5, 7.4 Hz, 1H), 4.04 (dd, *J* = 8.6, 5.2 Hz, 1H), 3.7 (s, 3H), 1.43 (s, 3H), 1.34 (s, 3H); **$^{13}$C-NMR (100 MHz, $CDCl_3$):** δ 171.7, 111.4, 74.1, 67.3, 52.4, 26.0, 25.6.

**D. Procedure for the conversion of methyl ester to *N*,*N*-dimethyl amide**

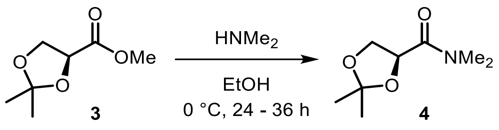


The amide **4** was prepared using a modified literature procedure(3). The acetonide protected methyl ester **3** (3.3 g, 20.6 mmol) was dissolved in 15 mL ethanol and cooled to 0 °C. Dimethyl amine was added in portion over two hours (5.0 equiv × 2). After stirring for 36 h, the reaction mixture was concentrated in vacuo and the residue was purified by column chromatography (50-60% ethyl acetate in petroleum ether) to obtain the amide **4** as a pale yellow liquid (2.5 g, 14.43 mmol, 70% yield); **$^1$H-NMR (400 MHz, $CDCl_3$):** δ 4.47 (t, *J* = 6.6 Hz, 1H), 4.08 (dd, *J* = 8.3, 6.4 Hz, 1H), 3.87 (dd,

$J$ = 8.1, 6.9 Hz, 1H), 2.88 (s, 3H), 2.7 (s, 3H), 1.15 (s, 6H); **$^{13}$C-NMR (100 MHz, $CDCl_3$):** δ 169.1, 111.6, 73.6, 66.5, 37.0, 35.9, 26.0, 25.9.

**E. Procedure for the conversion of amide to isopropenyl ketone (5)**

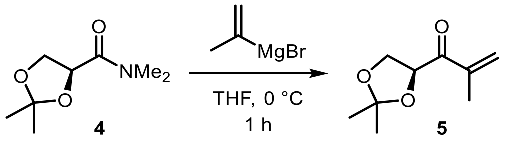


The isopropenyl ketone **5** was prepared using a modified literature procedure(3).In an oven dried 25 mL round bottom flask, the amide **4** was taken (1.0 g, 5.7 mmol) under argon along with dry THF. The resulting solution was cooled to 0 °C. Freshly prepared isopropenylmagnesium bromide was the added dropwise to the solution of **4** at 0 °C. The reaction mixture was stirred for 1 h at rt and then quenched with aqueous $NH_4Cl$ solution. The organic layer was extracted with diethyl ether. The combined organic layer was concentrated in vacuo and the residue was purified by column chromatography (7-10% ethyl acetate in petroleum ether) to obtain **5** as a pale yellow liquid (2.5 g, 14.43 mmol, 70% yield); **$^{1}$H-NMR (400 MHz, $CDCl_3$):** δ 6.00 (s, 1H), 5.89 (S, 1H), 5.05 (t, $J$ = 6.7 Hz, 1H), 4.20 (t, $J$ = 7.9, 1H), 4.05 (dd, $J$ = 7.9, 6.5 Hz, 1H), 1.88 (s, 3H), 1.39 (d, $J$ = 3.2 Hz, 6H); **$^{13}$C-NMR (100 MHz, $CDCl_3$):** δ 197.5, 142.6, 126.6, 110.6, 76.7, 66.3, 25.7, 25.4, 17.7.

**F. Procedure for the ozonolysis of isopropenyl ketone 5**

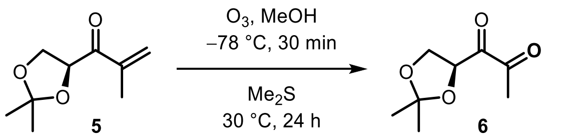


The diketone **6** was prepared using a modified literature procedure(4). The ketone **5** (0.7 mmol) was taken in a 25 mL round bottom flask along with 5 mL dry MeOH, and the solution was cooled at –78 °C for 10 minutes. Then $O_2$

was passed through this solution for 5 min followed by $O_3$ (at 40% power) until the starting consumed (monitored by TLC). $Me_2S$ (0.5 mL) was then added, and the solution was stirred at –78 °C for 10 min, followed by 24 h 30 °C. The reaction mixture was concentrated under reduced pressure and diluted with 3 mL water. The organic layer was extracted with ethyl acetate and washed with water. The combined organic layer was concentrated in vacuo and the residue was purified by column chromatography (20% ethyl acetate in petroleum ether) to obtain acetonide-protected DPD {**6**} as a yellow oil (70.0 mg, 0.406 mmol, 58% yield); **$^1$H-NMR (400 MHz, $CDCl_3$):** δ 5.10 (dd, *J* = 7.9, 5.4 Hz, 1H), 4.33 (t, *J* = 8.5 Hz, 1H), 3.96 (dd, *J* = 8.9, 5.4 Hz, 1H), 2.36 (s, 3H), 1.44 (s, 3H), 1.39 (s, 3H); **$^{13}$C-NMR (100 MHz, $CDCl_3$):** δ 198.1, 194.6, 111.3, 76.8, 66.0, 26.0, 25.3, 24.5.

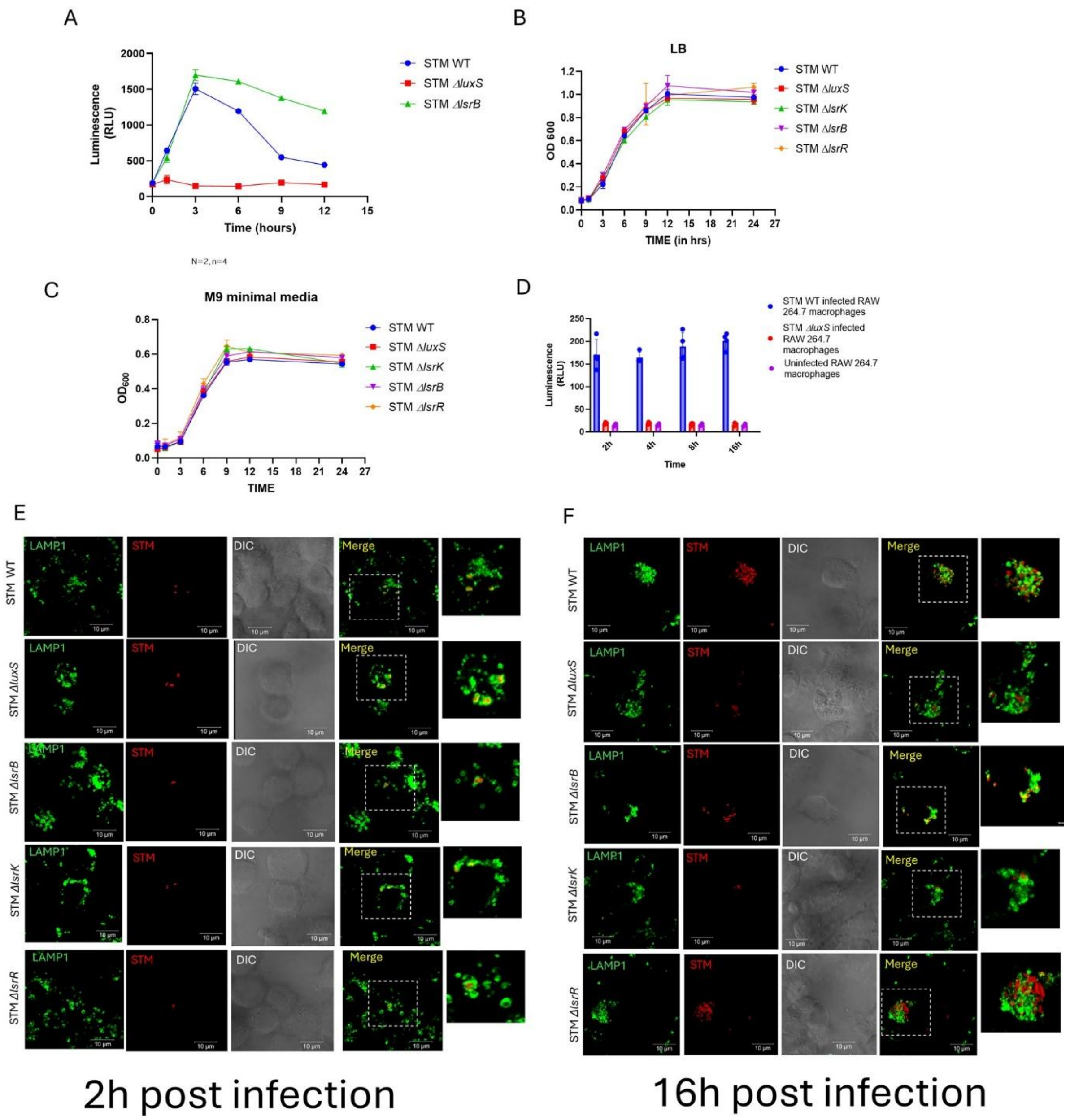


**Fig. S1 LuxS/AI-2 signaling is not required for *in vitro* growth but is essential for intracellular survival. (A)** STM WT, STM *ΔluxS,* and STM *ΔlsrB* AI-2 production in LB media. Represented as Mean+/-SD of N=3, n=4. Growth kinetics study upon deletion of gene *luxS*, *lsrB*, *lsrk*, and *lsrR* **(B)** LB media, **(C)** Minimal media. Represented as Mean+/-SD of N=3, n=3, **(D)** Autoinducer assay – AI-2 activity in RAW 264.7 macrophages upon infection**.**

Represented as Mean+/-SD of N=3, n=4. **(E)**and **(F)** Fluorescence microscopy of STM infection at 2h and 16h post-infection in RAW 264.7 macrophages. Representative of N=2, n>=10

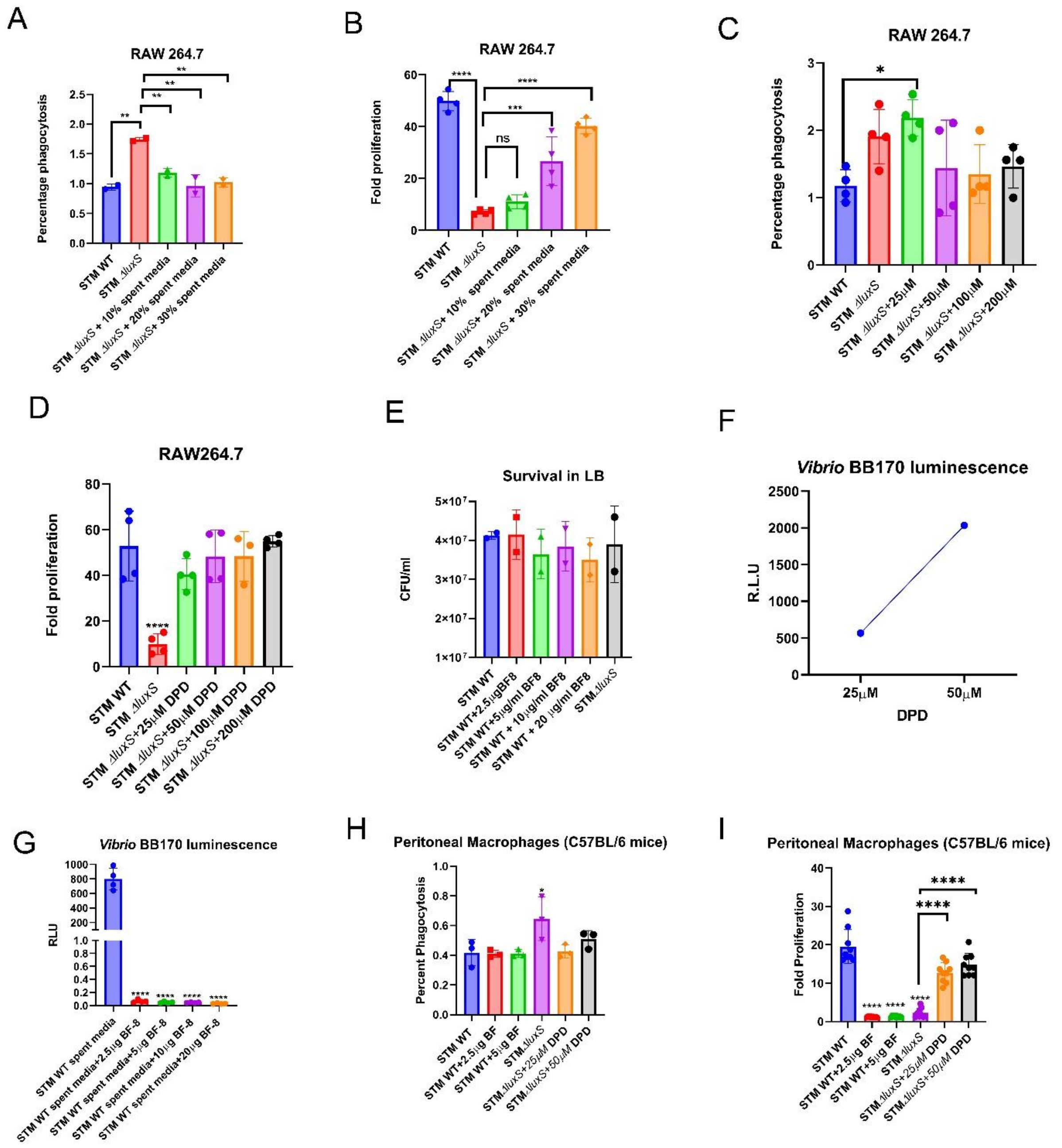


**Fig. S2 Spent media or exogenous AI-2 enhances STM survival in macrophages, while inhibition of AI-2 signaling attenuates it. (A)**

Percentage phagocytosis **(B)** Fold proliferation in RAW 264.7 macrophages of STM *ΔluxS* upon treatment of STM WT spent media at 10%, 20%, and 30%. Represented as Mean+/-SD of N=3, n=3. **(C)**Percentage phagocytosis, **(D)** Fold proliferation of STM in RAW 264.7 macrophages upon treatment of synthetic DPD molecule. Represented as Mean+/-SD of N=3, n=3. **(E)** Survival assay of STM WT and STM *ΔluxS* in the presence of inhibitor BF-8. Represented as Mean+/-SD of N=2. **(F)** Autoinducer assay- Light production by *Vibrio* BB170 for synthetic DPD molecule. Represented as Mean+/-SD of N=2, n=3. **(G)** Luminescence by *Vibrio* BB170 in STM WT spent media (3h growth spent media) with or without inhibitor BF-8. Represented as Mean+/-SD of N=2, n=3. **(H)**Percentage phagocytosis, **(I)** Fold proliferation of STM in peritoneal macrophages upon treatment of DPD and BF-8 inhibitor. Represented as Mean+/-SD of N=3, n=3. One-way ANOVA with Dunnet's post-hoc test was used to analyze the data; p values **** $p < 0.0001$, *** $p < 0.001$, ** $p<0.01$, * $p<0.05$.

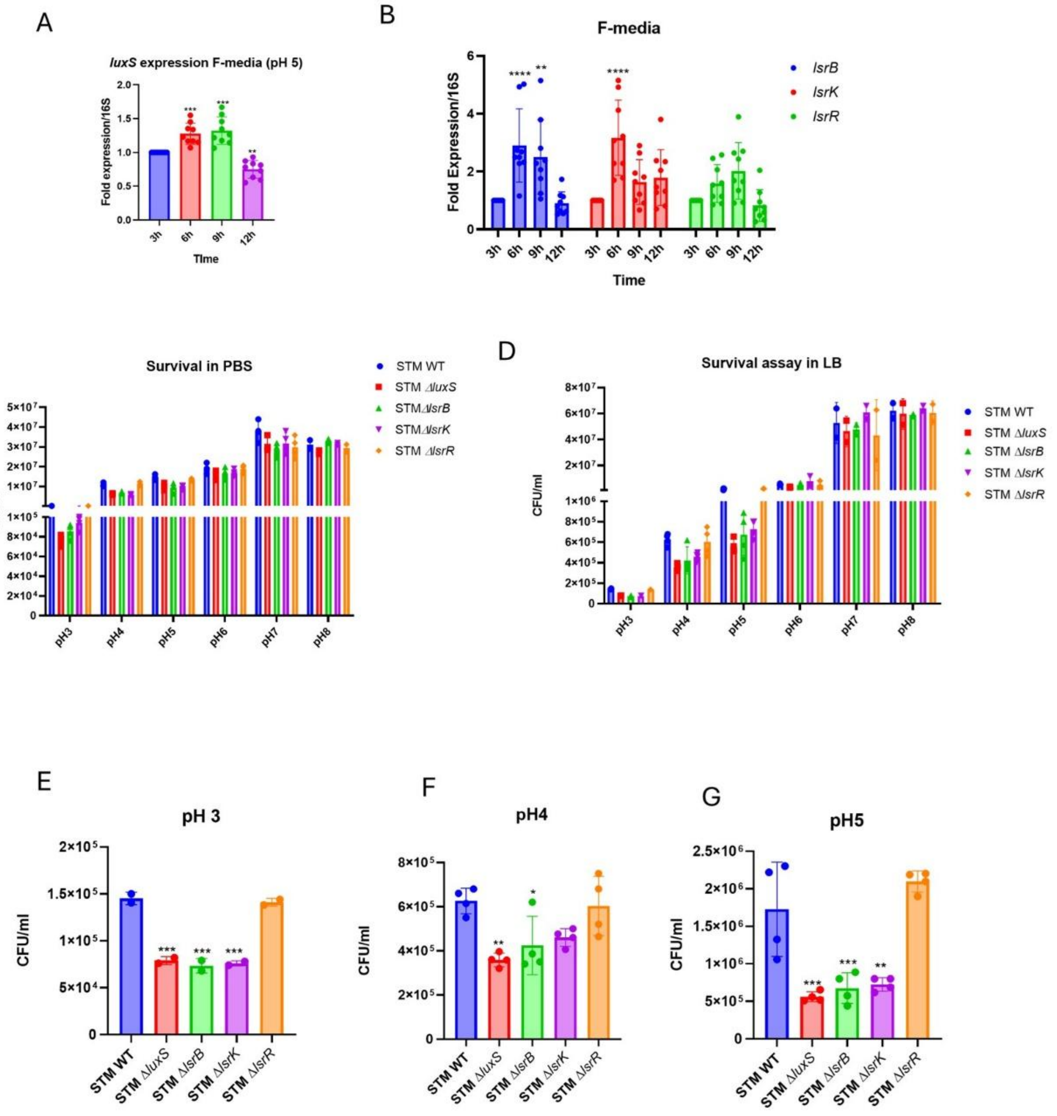


**Fig. S3 AI-2 signaling is induced in an acidic environment and increases survival at acidic pH. (A)** *luxS* **(B)** *lsr* operon gene *lsrB*, *lsrK* and *lsrR* expression study in STM WT upon growth in F-media (pH 5). Represented as Mean+/-SD of N=3, n=3. STM WT, STM *ΔluxS*, STM *ΔlsrB*, STM *ΔlsrK,* and STM *ΔlsrR* survival in**(C)** PBS and, **(D)** LB media with different range of pH (3-8). **(E)** Survival at pH 3 LB media **(F)** Survival at pH 4 LB media **(G)** Survival at pH 5 LB media. One-way ANOVA with Dunnet's post-hoc test was used to analyze the data; p values **** $p < 0.0001$, *** $p < 0.001$, ** $p<0.01$, * $p<0.05$. Two-way Anova was used to analyze the grouped data; p values **** $p < 0.0001$, *** $p < 0.001$, ** $p<0.01$, * $p<0.05$

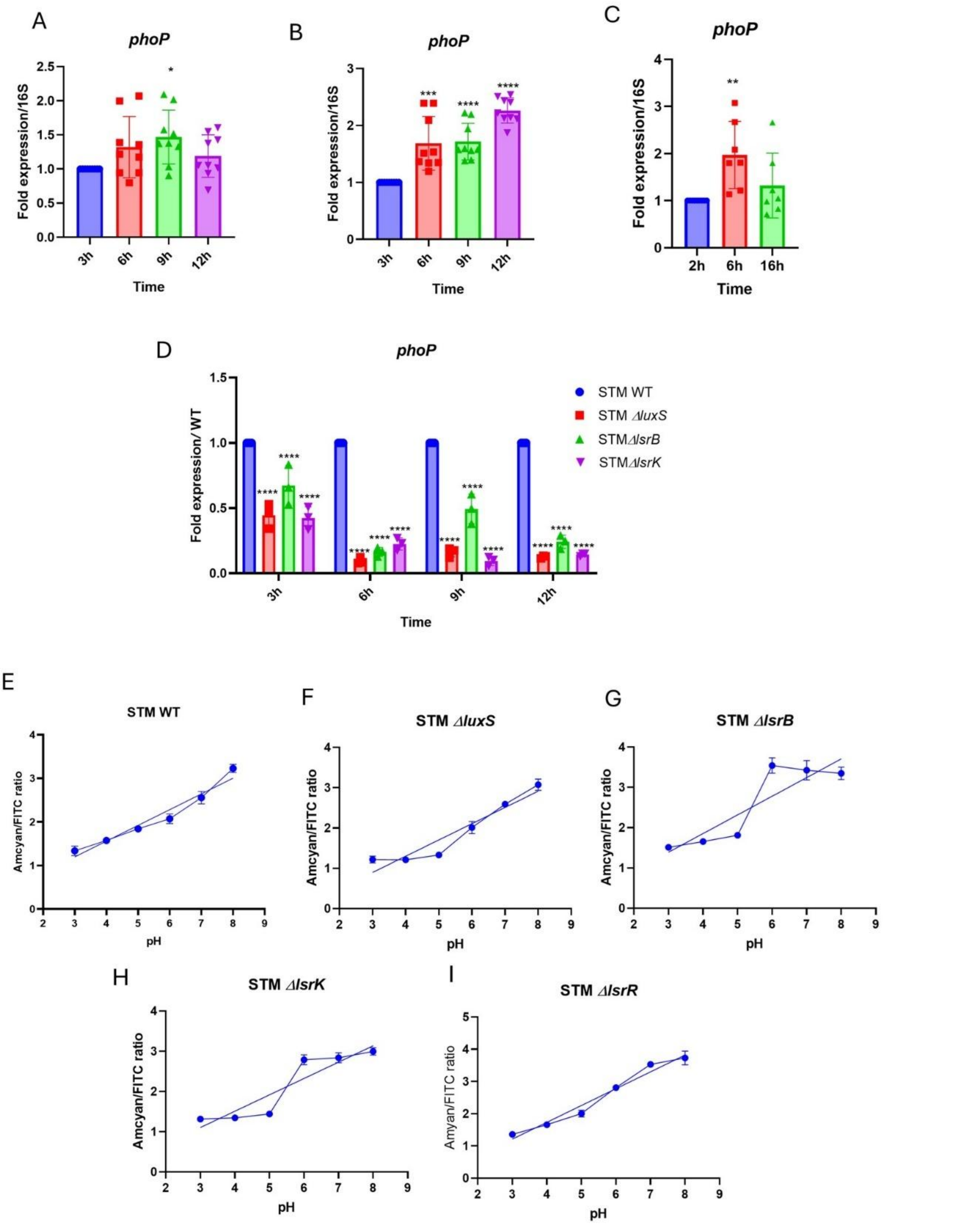


**Fig. S4 *phoP* expression is regulated by LuxS/AI-2 signaling and helps in maintaining the cytosolic pH.** mRNA expression of *phoP* gene in STM WT **(A)** LB media **(B)** F-media (pH 5), **(C)** Upon infection in RAW 264.7 macrophages. Represented as Mean+/-SD of N=3, n=3, **(D)** *phoP* gene expression in *STM* WT, STM *ΔluxS,* STM *ΔlsrB,* and STM *ΔlsrK,* in F-media (pH 5). Represented as Mean+/-SD of N=3, n=3. Standard curve of amcyan/FITC ratio in the range of pH 3-8 PBS of phuji plasmid containing STM strains **(E)** STM WT, **(F)** STM *ΔluxS*, **(G)** STM *ΔlsrB*, **(H)** STM *ΔlsrK, and* **(I)** STM *ΔlsrR*. One-way ANOVA with Dunnet's post-hoc test was used to analyze the data; p values **** $p < 0.0001$, *** $p < 0.001$, ** $p<0.01$, * $p<0.05$. Two-way Anova was used to analyze the grouped data; p values **** $p < 0.0001$, *** $p < 0.001$, ** $p<0.01$, * $p<0.05$

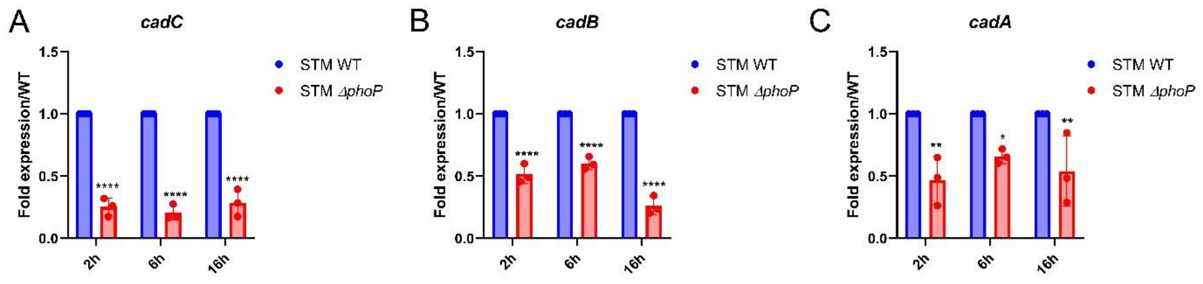


**Fig. S5 PhoP regulates *cadBC/A* genes to maintain cytosolic pH.** mRNA expression of **(A)***cadA,* **(B)** *cadB,* and **(C)** *cadC* gene in STM WT and STM Δ*phoP upon* infection in RAW 264.7 macrophages. Represented as Mean+/- SD of N=2, n=3. Two-way Anova was used to analyze the grouped data; p values **** p < 0.0001, *** p < 0.001, ** p<0.01, * p<0.05.

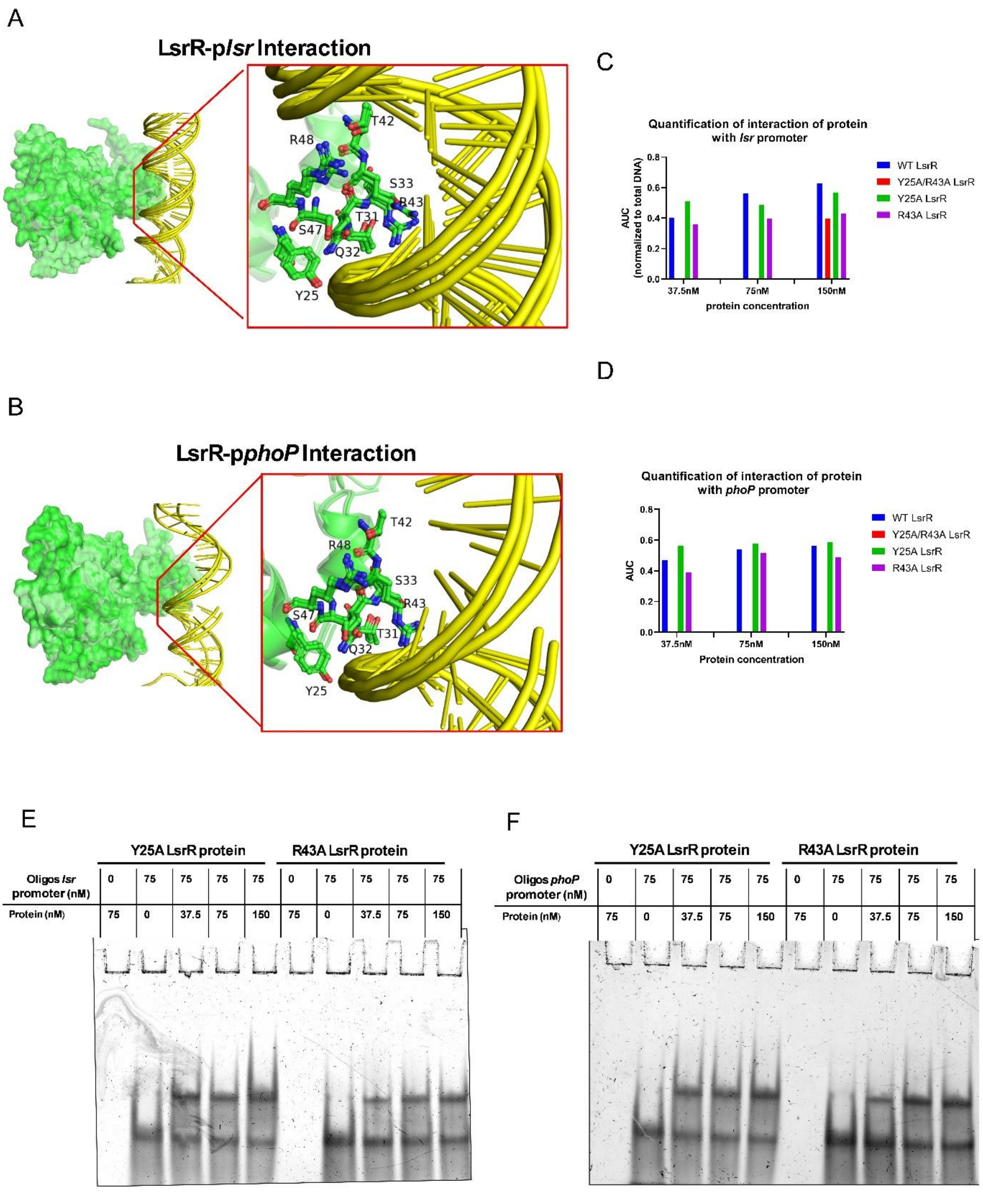


**Fig. S6 LsrR controls *phoP* expression by binding to its promoter. (A,B)** Superimposed close-up structural models showcasing key interactions between LsrR (green) residues and the *phoP* and *lsr* promoters (yellow), emphasizing critical contact points. Three individual AlphaFold3 structural models with the highest prediction rankings were aligned, and the interacting residues are displayed in a ball-and-stick representation. **(C, D)** Quantification of representative gel image of Fig. 4C, D, and extended Fig. 6E, F. Quantification done as the area under the curve (AUC) of protein-DNA bound/ (AUC protein DNA bound +AUC protein Unbound DNA)**.** Electrophoretic Mobility Shift Assay (EMSA) Y25A LsrR and R43A LsrR protein with double-stranded **(E)** *lsr* promoter and **(F)** *phoP* promoter.

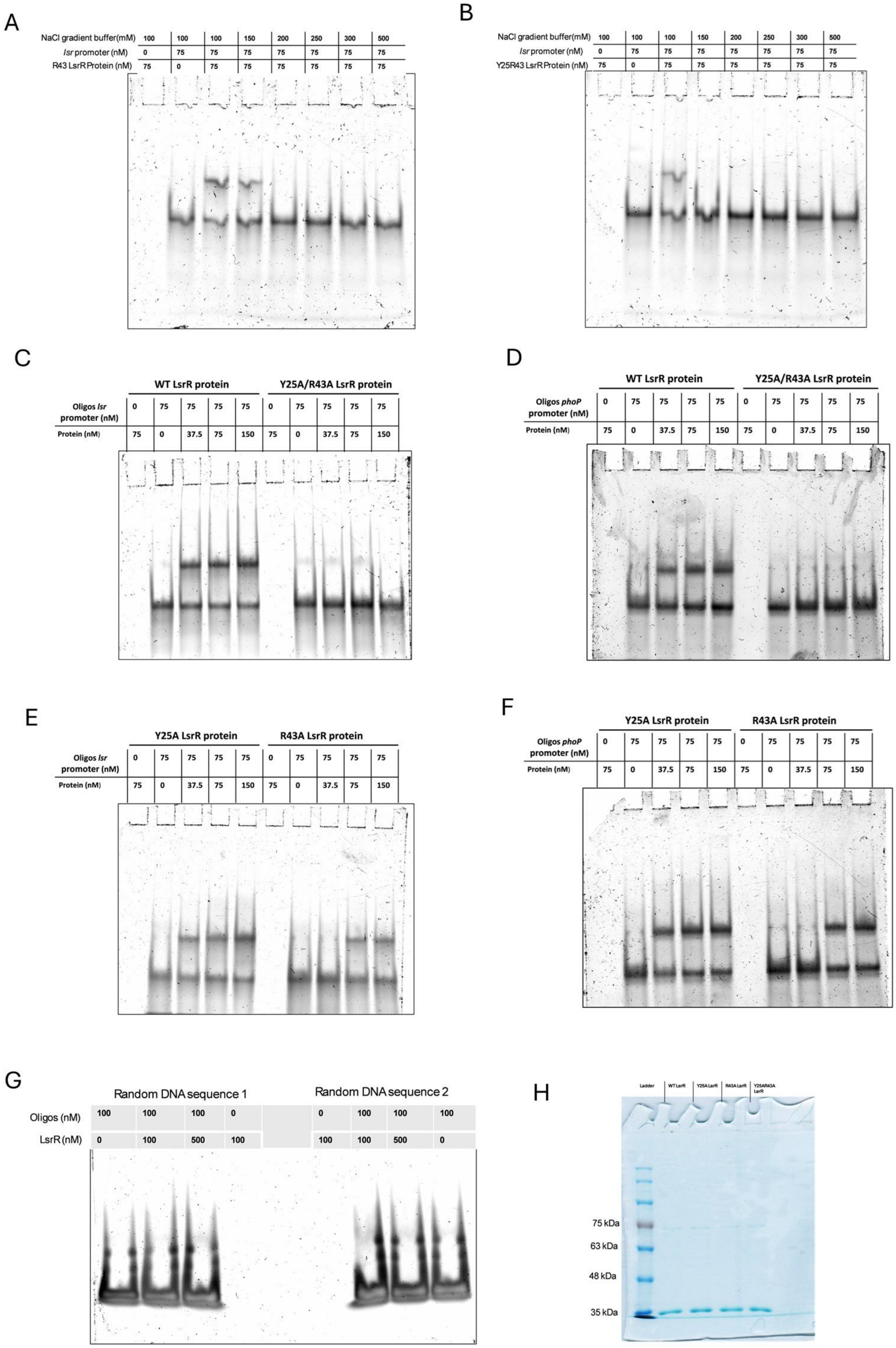
A
NaCl gradient buffer(mM) 100 100 100 150 200 250 300 500
lsr promoter (nM) 0 75 75 75 75 75 75 75
R43 LsrR Protein (nM) 75 0 75 75 75 75 75 75
B
NaCl gradient buffer(mM) 100 100 100 150 200 250 300 500
lsr promoter (nM) 0 75 75 75 75 75 75 75
Y25R43 LsrR Protein (nM) 75 0 75 75 75 75 75 75
C
WT LsrR protein
Y25A/R43A LsrR protein
Oligos lsr promoter (nM) 0 75 75 75 75 0 75 75 75 75
Protein (nM) 75 0 37.5 75 150 75 0 37.5 75 150
D
WT LsrR protein
Y25A/R43A LsrR protein
Oligos phoP promoter (nM) 0 75 75 75 75 0 75 75 75 75
Protein (nM) 75 0 37.5 75 150 75 0 37.5 75 150
E
Y25A LsrR protein
R43A LsrR protein
Oligos lsr promoter (nM) 0 75 75 75 75 0 75 75 75 75
Protein (nM) 75 0 37.5 75 150 75 0 37.5 75 150
F
Y25A LsrR protein
R43A LsrR protein
Oligos phoP promoter (nM) 0 75 75 75 75 0 75 75 75 75
Protein (nM) 75 0 37.5 75 150 75 0 37.5 75 150
G
Random DNA sequence 1
Random DNA sequence 2
Oligos (nM) 100 100 100 0 0 100 100 100
LsrR (nM) 0 100 500 100 100 100 500 0
H
Ladder
WT LsrR
Y25A LsrR
R43A LsrR
Y25AR43A LsrR
75 kDa
63 kDa
48 kDa
35 kDa

**Fig. S7 LsrR interacts with the *lsr* and *phoP* promoters through the Y25 and R43 amino acid residues.** Electrophoretic Mobility Shift Assay (EMSA) of *lsr* promoter with **(A)** R43A LsrR and **(B)** Y25A/R43A LsrR protein in increasing concentrations of NaCl in the binding buffer. EMSA of WT LsrR, Y25A/R43 LsrR, Y25A LsrR, and R43A LsrR with single-stranded (90 bp) *lsr* promoter **(C,E)** and *phoP* promoter**(D,F)**. EMSA of WT LsrR protein with random 60 bp DNA sequence **(G)**. SDS-PAGE of purified protein (1ug protein of each calculated by Bradford assay, molecular weight of LsrR ~35kDa) **(H).**

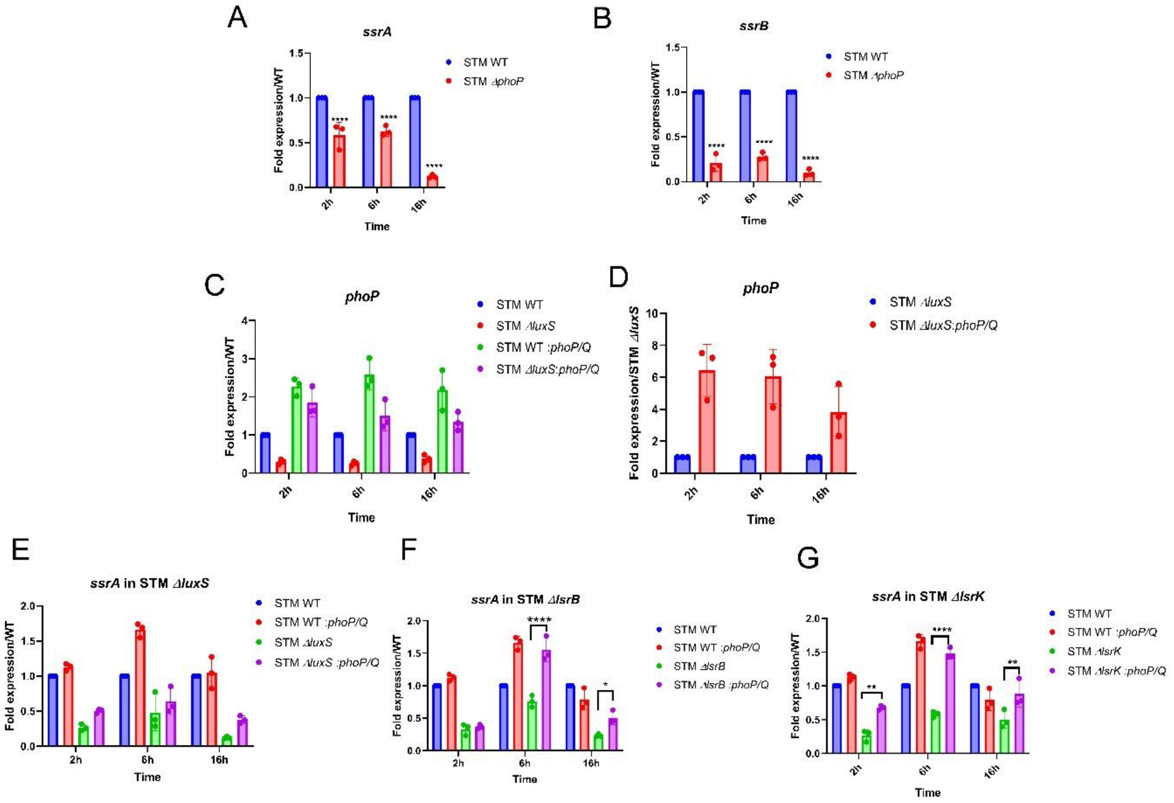

**Fig. S8 LuxS/AI-2 signaling regulates *ssrB/ssrA* gene expression through PhoP.** mRNA expression of **(A)***ssrA,* **(B)** *ssrB* gene in STM WT and STM Δ*phoP* *upon* infection in RAW 264.7 macrophages. Represented as Mean+/-SD of N=2, n=3. **(C) and (D)** mRNA expression of *phoP* gene in STM WT: pQE60-*phoP/phoQ, ΔluxS*: pQE60-*phoP/phoQ* upon infection in RAW 264.7 macrophages. Represented as Mean+/-SD of N=3, n=3. *ssrA* gene expression in *phoP/phoQ* cloned strain **(E)** STM *ΔluxS,* **(F)** STM *ΔlsrB,* **(G)** STM *ΔlsrK*. Represented as Mean+/-SD of N=3, n=3. One-way ANOVA with Dunnet's post-hoc test was used to analyze the data; p values **** $p < 0.0001$, *** $p < 0.001$, ** $p<0.01$, * $p<0.05$. Two-way Anova was used to analyze the grouped data; p values **** $p < 0.0001$, *** $p < 0.001$, ** $p<0.01$, * $p<0.05$.

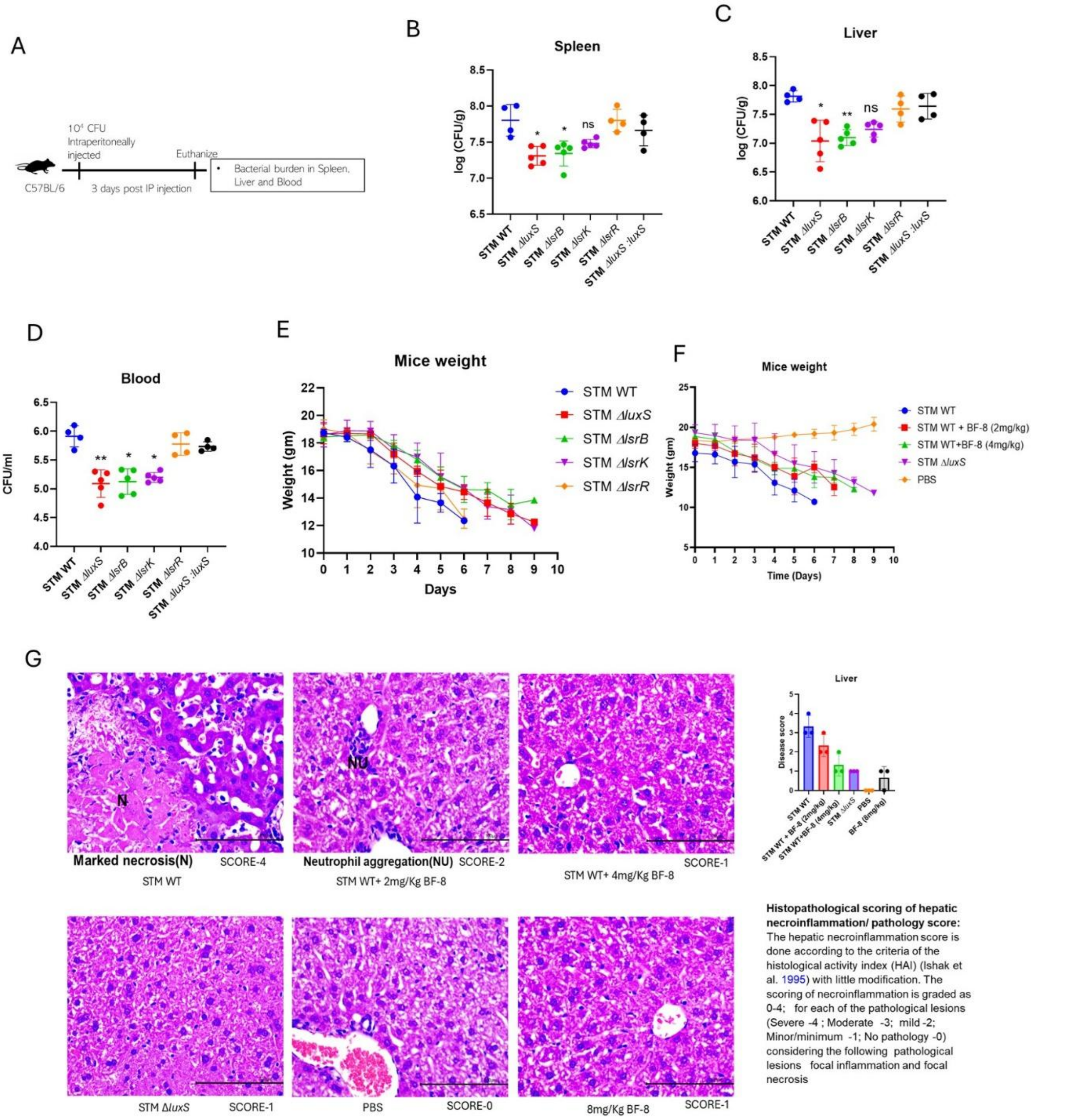


**Fig. S9 LuxS/AI-2 signaling facilitates colonization at secondary organ sites. (A)** The experimental protocol for organ burden in C57BL/6J mice by peritoneal infection of $10^4$ CFU per mouse. The organ burden post 3 days of peritoneal infection **(B)** Spleen, **(C)** Liver, and dissemination in **(D)** Blood. Represented as Mean+/-SD of N=2, n=5. **(E)** C57BL/6J mice infected by orally gavaging $10^8$ CFU per mouse. Mice weight reduction was noted on each day of post-infection. **(F)** C57BL/6J mice infected by orally gavaging $10^8$ CFU per mouse and BF-8 inhibitor treatment were given at alternative day of infection. Mice weight reduction was noted on each day of post-infection. Represented as Mean+/-SD of N=2, n=5. **(G)** The hematoxylin and eosin staining of the sections of the liver of C57Bl/6 mice infected by orally gavaging $10^7$ CFU per mouse and BF-8 inhibitor. treatment was given on the alternative day of infection. One-way ANOVA (Kruskal Wallis) with Dunn's post-hoc test was used to analyze organ burden in mice.

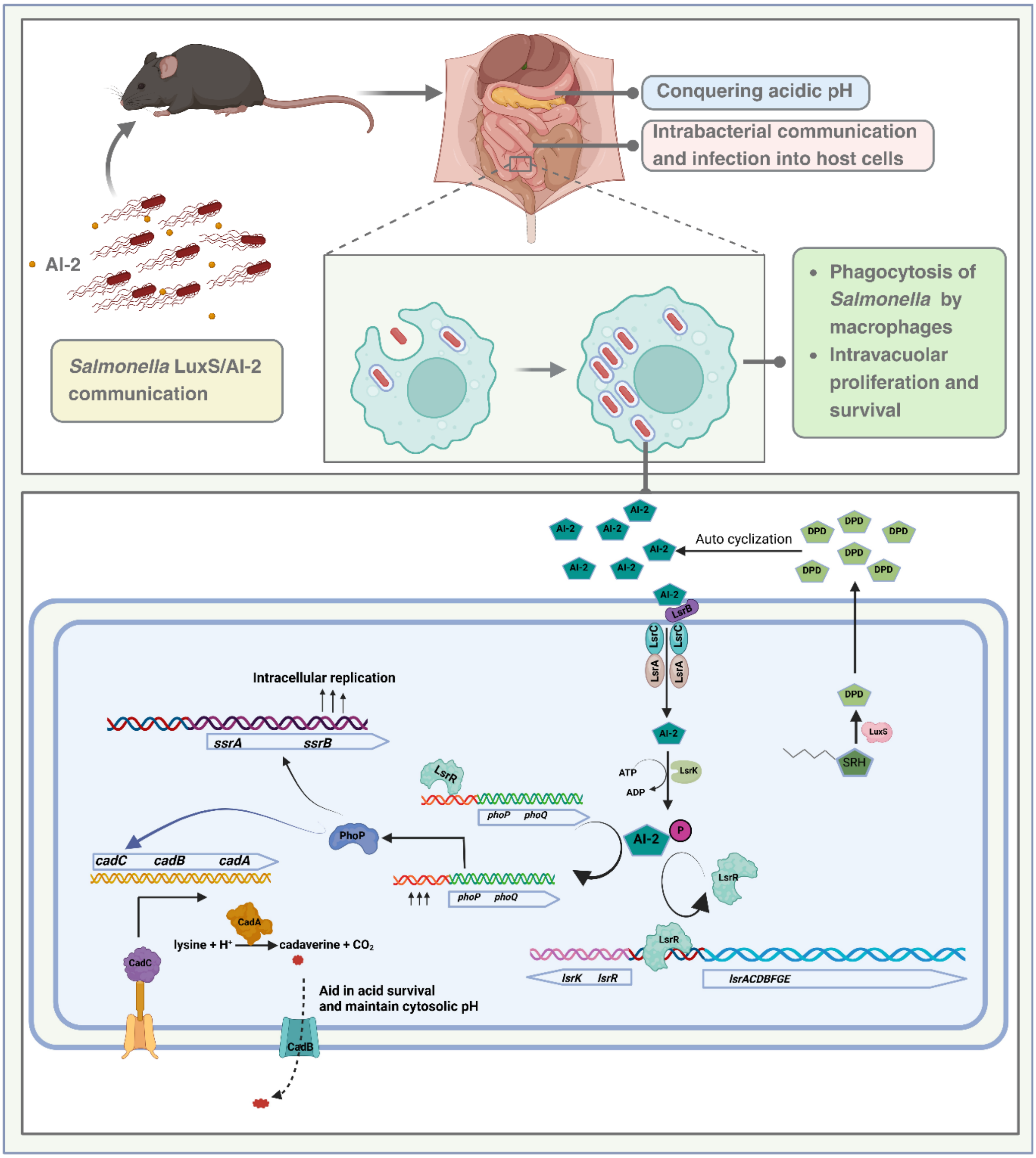


**Fig. 10 Graphical abstract**

**Table S1**

All the Bacterial strains and plasmids used in this study are given in Table -

| Strains/Plasmids | Characteristics | Sources |
|---|---|---|
| STM WT | No antibiotics | Gift from Prof. M. Hensel |
| STMΔ*luxS* | Chl$^{R}$ | This study |
| STMΔ*lsrB* | Kan$^{R}$ | This study |
| STMΔ*lsrK* | Chl$^{R}$ | This study |
| STMΔ*lsrR* | Kan$^{R}$ | This study |
| STMΔ*luxS* Δ*lsrR* | Chl$^{R}$ Kan$^{R}$ | This study |
| STMΔ*luxS:luxS* | Chl$^{R}$, Amp$^{R}$ (complemented in pQE60 plasmid) | This study |
| STMΔ*phoP* | Chl$^{R}$ | Laboratory stock |
| *Vibrio campbellii* ATTC BAA-1117 | No antibiotics | ATTC |
| pKD3 plasmid | Chl$^{R}$ resistance cassette | Laboratory stock |
| pKD4 plasmid | Kan$^{R}$ resistance cassette | Laboratory stock |

| | | |
|---|---|---|
| pKD46 plasmid | Plasmid expressing λ-red recombinase system, $Amp^R$ | Laboratory stock |
| pQE60 vector | Low copy number plasmid, $Amp^R$ | Laboratory stock |
| pQE60-*phoP*-*phoQ* Complemented plasmid | $Amp^R$ | This study |
| pBAD:pHuji | $Amp^R$ | Add gene |
| pET28a(+) | $Kan^R$ | Laboratory stock |
| pET28a(+)- *lsrR* | $Kan^R$ | This study |

**Table S3**

**Primers used in this study:**

| Primers | Sequence 5'.......3' |
|---|---|
| *luxS* knockout forward primer | CGGAGGTGACTAAATGCCATTATTAGATAGCTTCGCAGTC**CATATGAATATCCTCCTTAG** |
| *luxS* knockout reverse primer | CTGGAACCGCTTACAAATAAGACTAAATATGCAGTTCCTG**GTGTAGGCTGGAGCTGCTTC** |
| *luxS* knockout confirmation forward primer | GCAAAACACGCCTGACCCAA |
| *luxS* knockout confirmation reverse primer | CAATACACTCTGGCATCGTG |
| *luxS* expression forward primer | GCTCCAGAATATGACGGGCA |
| *luxS* expression reverse primer | CGCACCGGCTTTTACATGAG |
| *luxS* cloning forward primer | **CGCGGATCC**ATCGGAGGTGACTAAATGCC |
| *luxS* cloning reverse primer | **CCCAAGCTT**TGGAACCGCTTACAAATAAGAC |
| *lsrR* knockout forward primer | CAAAGTAAAGCCAGGTTATGACAATGAGCGATAATACGTTGG**CATATGAATATCCTCCTTAG** |
| *lsrR* knockout reverse primer | GAATTATTTTCCCTGCGGTTTTCTGATCGGTAACCAGTGC**GTGTAGGCTGGAGCTGCTTC** |
| *lsrR* knockout confirmation forward primer | GTGGCGTTAATCACGGTTAT |

| | |
|---|---|
| *lsrR* knockout confirmation reverse primer | TATTGAATTGAGGTAAGTGT |
| *lsrR* expression forward primer | GAATATCGCCTACTGCGCCT |
| *lsrR* expression reverse primer | AAATAGCGTGCGGGATGTGA |
| *lsrB* knock out forward primer | ATGGCAAGACACAGCATTAAAATGATCGCCTTACT CACTGCATATGAATATCCTCCTTAG |
| *lsrB* knock out reverse primer | TGAAAATGACACGCTCCGGCAATAACACAATGCC GTTACCGTGTAGGCTGGAGCTGCTTC |
| *lsrB* knockout confirmation forward primer | CGTCACCAAATCCTTGAATG |
| *lsrB* knockout confirmation reverse primer | CCACAGCCTTTTAATGCGAA |
| *lsrB* expression forward primer | AGAGTTTGGCCTGTGGGATG |
| *lsrB* expression reverse primer | GGTGAAACGGTGACTTTGCC |
| *lsrK* knock out forward primer | TGGCTCGACTCTGTACCCATACTGAATCAGGACATT ACCTCATATGAATATCCTCCTTAG |
| *lsrK* knock out reverse primer | CGCTTTCCAGAGGGATGTCGTTAAGCCACCATCG ACCAGGTGTAGGCTGGAGCTGCTTC |
| *lsrK* knockout confirmation forward primer | CGGTAACTATATCAATGCAC |
| *lsrK* knockout confirmation reverse primer | GCGGTATTCATGATTCTTCT |

| | |
|---|---|
| *lsrK* expression forward primer | GTGAAATGTGACCGAGCAGC |
| *lsrK* expression reverse primer | CTTTATGCTCAGCGGCGAAC |
| *phoP* expression forward primer | GATCTCTCACGCCGGGAATT |
| *phoP* expression reverse primer | TGACATCGTGCGGATACTGG |
| *phoQ* expression reverse primer | GCAGCAAACGAAAGGTGGTT |
| *phoQ* expression reverse primer | TTTGCTCGCCATTTTCTGCC |
| *phoP/phoQ* cloning forward primer | CGCGGATCCGAGATGATGCGCGTACTGGTT |
| *phoP/phoQ* cloning reverse primer | CCCAAGCTTTGGAAGAACGCACAGAAATGT |
| *ssrA* expression forward primer | ACCGCCCATCATTTTAGCCA |
| *ssrA* expression reverse primer | AACCGGAGGGATACGTCTGA |
| *ssrB* expression forward primer | CGGTGTGTTTCGACGGTTTT |
| *ssrB* expression reverse primer | ACGCTGACACGACCAATCAT |

| | |
|---|---|
| *ssaV* expression forward primer | CGCCGCAAAAAGTCTGTGGT |
| *ssaV* expression reverse primer | GGGACGCCGGTATCCTCAAA |
| *spiC* expression forward primer | ACCTAAGCCTTGTCTTGCCT |
| *spiC* expression reverse primer | CCATCCGCTGTGAGCTGTAT |
| *cadA* expression forward primer | CTGAACCTGCGCGTCAAAAA |
| *cadA* expression reverse primer | TTTCGGCAGCACTTCAAACG |
| *cadB* expression forward primer | TTGGCCTATTCCTGGTGCTG |
| *cadB* expression reverse primer | GCGTTTCGGGTTTTTCACCA |
| *cadC* expression forward primer | TCAATCAGCGCCACTATCGG |
| *cadC* expression reverse primer | GATTAGGCCAGGGCTGTGTT |
| *lsrR* cloning forward primer | CATGCCATGGACAATGAGCGATAATACGTTG |
| *lsrR* cloning reverse primer | CCGCTCGAGTTTTTCAATAATTTGAATTATTTTCCCTGC |

| | |
|---|---|
| Y25A *lsrR* SDM forward primer | CGTATTGCCTGGTTCGCCTATCACGATGG |
| Y25A *lsrR* SDM reverse primer | CCATCGTGATAGGCGAACCAGGCAATACG |
| R43A lsrR SDM forward primer | CTGGGGCTAACCGCGCTAAAGGTTTCTCG |
| R43A lsrR SDM reverse primer | CGAGAAACCTTTAGCGCGGTTAGCCCCAG |
| *lsr* promoter sequence | GACCAAATAACTACTACCGTTTTGAACAATTTCTTTT TCAAAAAACATTTGTTCAGTCCCGTCAGTCAACATT GAGGGAGCGGAGGCAAC |
| *phoP* promoter sequence | AAGAGGGTGACTATTTGTCTGGTTTATTAACTGTTTAT CCCCAAAGCACCATAATCAACGCTAGACTGTTCTT ATTGTTAACACAAGGGA |
| 16s rRNA forward primer | CGCTTCTCTTTGTATGCGCC |
| 16s rRNA reverse primer | TCGTCAGCTCGTGTTGTGAA |

**Table S2**

**Media Composition**

| | |
|---|---|
| **M9 Minimal Media**: | 2mM Magnesium Sulphate, 0.1mM Calcium Chloride, 0.4% Glucose, 10X M9 salts [Di-Sodium hydrogen phosphate (64g in 1000ml), Potassium Di-hydrogen phosphate (15g in 1000ml), Sodium Chloride(2.5g in 1000ml), Ammonium Chloride(5g in 1000ml)] at a final concentration of 1X. |
| **F-Media:** | 8μM Magnesium Chloride, 38mM Glycerol, 0.1% Cas-amino acids, 10X F-media salts (50mM Potassium Chloride, 75mM Ammonium Sulphate, 5mM Potassium Sulphate, 10mM Potassium Di-hydrogen phosphate, 1M Bis-Tris) at final concentration of 1X. Final pH adjusted to 5. |
